# Are Electrical Characterizations Consistent with the Cytochrome Structures of *Geobacter* ‘Nanowires’

**DOI:** 10.1101/2023.10.10.561676

**Authors:** Matthew J. Guberman-Pfeffer

## Abstract

Electrically conductive filaments from *Geobacter sulfurreducens* were reported to be pili with metallic-like conductivity, and yet were later shown to be redox-active cytochromes by cryogenic electron microscopy. It has recently been argued that the filaments were simply misidentified, implying that key observations formerly used to refute the involvement of cytochromes in conductivity now must be ascribed to them. Herein, the temperature, pH, voltage, crystallinity, charge propagation, and aromatic density-related dependencies of the conductivity reported for putative pili are re-examined in light of the CryoEM structures of cytochrome filaments. It is demonstrated that:

1. Electrons flow through cytochrome filaments in a succession of redox reactions for which the energetics are physically constrained and the kinetics are largely independent of protein identity for highly conserved heme packing geometries. Computed heme-to-heme electron transfer rates in cytochrome filaments agree, on average, within a factor of 10 of rates experimentally determined in other multi-heme proteins with the same heme packing geometries.
2. T-stacked heme pairs, which comprise nearly or exactly half of all heme pairs in cytochrome filaments are electronic coupling-constrained bottlenecks for electron transfer that set the rate-limiting reaction to the µs timescale, which is *fast enough* compared to typical ms enzymatic turnover. Tuning the conductivity of cytochromes over the reported ∼10^7^-fold range for filaments from *G. sulfurreducens* strains with pili variants seems both physically implausible and physiologically irrelevant if those filaments are supposed to be cytochromes.
3. The protein-limited flux for redox conduction through a 300-nm filament of T- and slip-stacked heme pairs is predicted to be ∼0.1 pA; a *G. sulfurreducens* cell discharging ∼1 pA/s would need at least 10 filaments, which is consistent with experimental estimates of filament abundance. The experimental currents for the Omc- S and Z filaments at a physiologically relevant 0.1 V bias, however, are ∼10 pA and ∼10 nA, respectively. Some of the discrepancy is attributable to the experimental conditions of a dehydrated protein adsorbed on a bear Au- electrode that contacts ∼10^2^ hemes, and in the case of conducting probe atomic force microscopy, is crushed under forces known to deform and change the electron transport mechanism through more highly-structured proteins.
4. Previously observed hallmarks of synthetic organic metallic-like conductivity ascribed to pili are inconsistent with the structurally resolved cytochrome filaments under physiological conditions, including (I) increased crystallinity promoting electron delocalization, (II) carbon nanotube-like charge propagation, and (III) an exponential increase-then-decrease in conductivity upon cooling, which was only explain by a model predicted on redox potentials known to be experimentally false. Furthermore, spectroscopic structural characterizations of OmcZ that attest to a huge acid-induced transition to a more crystalline state enhancing conductivity either strongly disagree with CryoEM analyses at higher pH values or give inconclusive results that can be overly interpreted.

Overall, a significant discrepancy currently exists—*not between theory and experiment*—but between the CryoEM cytochrome filament structure in one hand and the other functional characterizations of *Geobacter* ‘nanowires’ in the other. The CryoEM structures, theoretical models, biological experiments, and kinetic analyses are all in agreement about the nature and rate of electron transfer in multi-heme architectures under physiological conditions, and stand opposed to the solid-state functional characterizations of *Geobacter* filaments reported to date. The physiological relevance and/or physical plausibility of some experiments should be examined further.

## 1. Introduction

Filaments from *Geobacter sulfurreducens* were reported nearly 20 years ago to be electrically conductive,^1^ and yet, an intense debate persists over their identity, structure, and *in vivo* mechanism.^2-5^ Two hypotheses have divided the field (Figure 1):^6^ (1) The filament is a supramolecular assembly of cytochromes that transfers electrons through a ‘bucket-brigade’ succession of reduction-oxidation (redox) reactions;^7, 8^ or (2) the filament is a supramolecular assembly of PilA proteins that delocalizes electrons through a crystalloid array of stacked aromatic residues to realize metallic-like conductivity.^9, 10^

**Figure 1.**
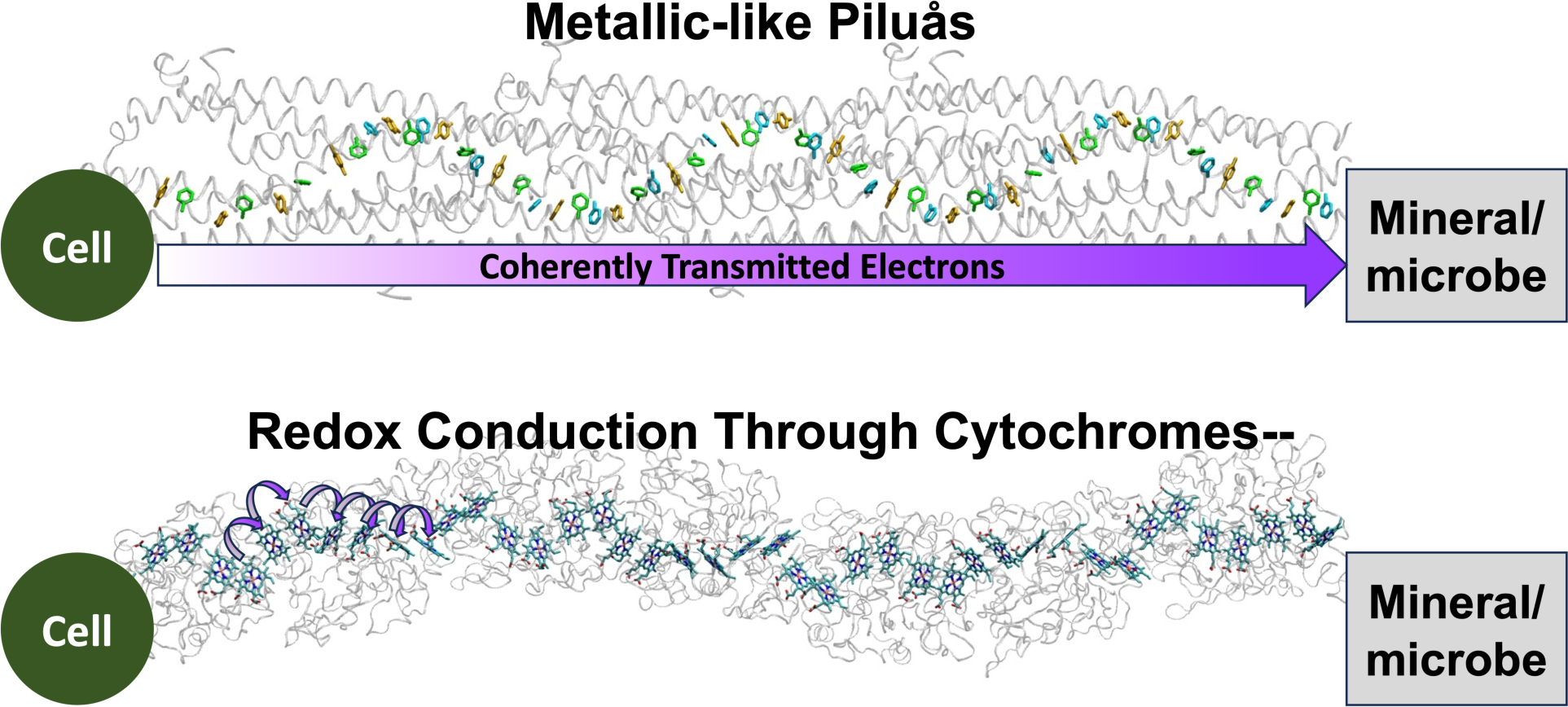
Schematics of the metallic-like pilus (MLP) and redox conduction through cytochrome (RCTC) hypotheses. The model of the pilus comes from Ref. 11. The structure of the cytochrome comes from Ref. 3.

Practitioners in the field have thought that the debate would be settled once the cryogenic electron microscopy (CryoEM) structure of the filament was resolved. Indeed, Malvankar and co-workers^5^ have likened the situation without CryoEM to the Indian parable of blind men who first encounter an elephant, whereupon each comes to a different conclusion about the animal’s identity from their limited perspective: Microbiologists thought of pili as electrical conduits to monomeric cytochromes that served as terminal reductases; electrochemists thought of pili as structural scaffolds for the dynamic assembly of cytochromes; and “[b]iophysicists, like us,” according to Malvankar and co-workers, “found that the filaments are intrinsically conductive…but did not know the mechanism.”^5^ Malvankar and co-workers concluded that “Lack of nanowire structure was everybody’s elephant in the room.”^5^

This telling of the history does not complete the analogy. It omits the parallel between the dispute among the blind men that erupted when some—in this case, prominently Malvankar and co-workers—concluded with absolute certainty (see below) that their mechanistic hypothesis was correct from an impressive and extensive assortment of functional and preliminary structural analyses^9-22^ in favor of the metallic-like pilus hypothesis.

There are now six CryoEM filaments, five of which support the redox conduction through cytochrome (RCTC) hypothesis.^3, 23-27^ The sixth filament structure is of the *G. sulfurreducens* pilus^28^ and shows that the metallic-like pilus (MLP) hypothesis—or any other aromatic-based conductivity—is completely impossible. And yet, the debate continues, perhaps because the absence of a CryoEM structure of the metallic-like pilus does not *prove* it cannot or does not exist. What seems more surprising is that data used to advocate for the MLP hypothesis^9, 17, 22^ is now claimed as consistent with the RCTC hypothesis.^3, 29, 30^ A re-examination of the history and assumptions in the field therefore seems warranted to set the stage for the questions examined by the present study. After the present author attended back-to-back talks that exclusively discussed either conductive pili or conductive cytochrome filaments from *G. sulfurreducens*, he overheard a prominent theorist exclaim “What is a poor theorist to think!” This article is the present author’s reply.

### 1.1. Simpler Explanations are Usually Right

The RCTC hypothesis was consistent with the long-established theory and function of redox cofactor chains in biology that connect catalytic centers.^31-38^ The hypothesis was supported by the prevalence of extracellular cytochromes expressed by *G. sulfurreducens*, as well as a wealth of spectroscopic and electrochemical data consistent with redox chemistry.^7, 8, 39-53^ It was also considered implausible that crystalline-like order could be maintained among rotatable aromatic sidechains throughout a non-covalent protein assembly at 300 K.^7, 54, 55^

The MLP hypothesis was inconsistent with what was known about biological electron transfer,^31^ but so too, it seemed, was the micron-length scale of electron transfer through the filaments.^56^ The MLP hypothesis came to be prominently advocated by Malvankar and colleagues as a “new paradigm for biological electron transfer and bioelectronics.”^10^ Malvankar and co-workers performed an extensive series of experiments that both excluded cytochromes from being involved in the conductivity of *G. sulfurreducens* biofilms and filaments,^13^ and characterized several properties inconsistent with redox conduction through cytochromes.^9, 11, 12, 14-22^ These experiments showed temperature,^9, 15^ pH,^9, 15, 19^ voltage,^9, 13^ crystallinity,^11, 17^ charge propigation,^15, 16^ and aromatic density-related^14, 18-20^ dependencies of the conductivity—sometimes under cytochrome denaturing or inhibiting conditions—similar to synthetic organic metals.^10^

Some of the observed hallmarks of metallic-like conductivity were debated on grounds of inappropriate experimental design.^12, 21, 39, 44, 57^ Other results were not reproduced in magnitude^58^ or sign^59^ by others, or the results were shown to depend heavily on experimental conditions that had not been controlled.^45^ Even with these issues, the MLP hypothesis was not refuted because, as Malvankar and co-workers argued, the hypothesis was supported “from multiple approaches, with different inherent assumptions.”^13^

Some theoretical work^60-62^—including two reports of homology modeling on which Malvankar was a co-author^11, 17^—provided support for the picture of a seamlessly stacked aromatic array for pilus conductivity. Other modeling efforts were skeptical^6, 7, 46, 54, 55, 63-66^ and dismissed out-of-hand by Malvankar and co-workers: “Although several models … have questioned whether sufficient pi-pi stacking of aromatic amino acids for metallic-like conductivity is possible, actual experimental results trump modelling, which can be based on faulty assumptions.”^16^

Malvankar and co-workers explained why some models were more useful than others: “No model can prove or disprove experimental results, but it is necessary for a model to be consistent with experimental observations.”^11^ This view implies that only models that validate experiments are acceptable, and that theory has no role in saying what is physically plausible—the present author and study herein disagrees. It is useful to recall that one of the earliest supports for the Periodic Table of the Elements came from Lecoq de Boisbaudran repeating his measurements on Gallium because Dmitri Mendeleev insisted that the experiments, not his predictions, were wrong; Dmitri was right; the experimentalist had made mistakes.^67^

But there was strong evidence for believing cytochromes could not account for the conductivity of *G. sulfurreducens* biofilms and filaments, as expressed by Malvankar and co-workers: “Long-range electron transport through biofilms via *c*-type cytochromes is physically impossible because the density of cytochromes is too low and cytochromes are spaced too far apart for electron hopping/tunneling through the biofilm.”^13^ And “the finding that denaturing cytochromes in pili preparations had no impact on conductivity definitively rules out a role for OmcS, or any other *c*- type cytochrome, in electron conduction along the pili.”^12^

The debate, however, continued: “In spite of these results the hypothesis [from others] that cytochromes are responsible for long-range conduction of electrons through biofilms and along the pili of *G. sulfurreducens* persists.”^13^ And yet, Malvankar and co-workers were among the first^3,23^ to publish a CryoEM-resolved structure of the outer-membrane cytochrome type S (OmcS) filament from *G. sulfurreducens*. Their article stressed the novelty of a cytochrome filament, while not discussing how the advocates of cytochrome-based conduction had been correct, or how “[t]he original hypothesis that OmcS makes up the filament”^68^ was considered over a decade earlier by contemporaries of Malvankar in the Lovely laboratory.^68-70^ Malvankar and co-workers concluded that “these [OmcS] nanowires are the same filaments that were previously thought to be type IV pili.”^3^ Apparently there was never any metallic-like pilus!

The simple and expected explanation had won (it seemed) over the fantastical vision of a “carbon nanotube-like”^16^ pilus. It is remarkable and commendable how willing Malvankar and co-workers were to change their opinion on the pili-versus-cytochrome dispute once faced with the CryoEM evidence of a cytochrome. The problem of interest here, however, is that the characterization data presented by Malvankar and co-workers in favor of the MLP hypothesis is *not* a matter of opinion. It is incumbent on us to ask: If that change of opinion is warranted, how was an entire edifice for a “new paradigm”^10^ of metallic-like conduction in biology predicated on a falsely identified protein? And, how can a suite of structural and electrical characterizations inconsistent with redox conduction through cytochromes, some of which were even performed under cytochrome denaturing or inhibiting conditions, now be attributed to cytochromes? Are there new mechanisms of electrical conductivity in cytochromes for us to learn, and if so, are they biologically relevant? Alternatively, are different filaments being structurally and functionally characterized? Or, as previously charged,^44, 57^ were the characterizations supporting metallic-like pili simply performed incorrectly: “To date [2015], the only results inconsistent with redox conduction [through cytochromes] … are those reported by Malvankar *et al.,* which have been addressed elsewhere and which have not been corroborated by others.”^8^ These are the questions that motivated the present study.

### 1.2. Blindness is a Choice of Our Assumptions

Malvankar and co-workers have already tried to resolve the cognitive dissidence by explaining: “There has never been any direct evidence that conductive *Geobacter* extracellular filaments are composed of PilA,”^3^ and “conduction along the length of a single *bona fide* PilA filament has not been demonstrated.”^3^ But why then did an article titled *Conductivity of Individual Geobacter Pili* by Malvankar and co-workers state three years earlier, “The estimates of individual *G. sulfurreducens* pilus conductivity reported here provide a key piece of data that is needed for assessing the diverse proposed models for the conductivity of *G. sulfurreducens* pili.”^18^ Note the absence of the term “putative” or any other qualifier to signal that the identity of the protein was a questionable interpretation instead of a proven fact. What was the basis for this misplaced confidence in the identity of the filament as a pilus?

Malvankar and co-workers argued at the time that immunogold labeling suggested, and atomic force microscopy (AFM) confirmed that cytochrome-like globules were aligned along pili.^13^ The spacing between those globules was >100 nm, too large for cytochromes to be involved in conductivity.^13^ The pili filament and the cytochrome-like globules were identified by their height profiles in AFM (3–6 vs. 5–7 nm),^13^ an approach still used by Malvankar and co-workers to help identify proteins before electrical characterization.^3, 29^

In the aftermath of the CryoEM structure, Malvankar and co-workers made it clear that the AFM and antibody labeling experiments gave the wrong result because researchers were blinded by their assumptions: “As cytochromes were not known to form filaments before our work, AFM images of filaments as well as these antibody-labeling results were interpreted as showing isolated OmcS monomers binding to the surfaces of the PilA filaments rather than showing antibodies directly binding to OmcS filaments.”^3^ However, redox chains in biology were well known at the time— even referred to as “molecular wire[s],”^32-37^—to routinely move electrons over long distances,^38^ and the “original hypothesis”^68^ was that the filaments were composed of OmcS. Also, contemporaries were well aware that “globular debris [on pili] may be aggregated pilins, cytochromes, or non-proteinaceous cell material.”^58^ To presume that those globules were in fact cytochromes from their height alone, and to argue that their >100 nm spacing on filaments—also presumed to be pili from height alone—”definitively rules out”^12^ the cytochrome hypothesis was clearly an overzealous argument for the favored MLP hypothesis. It is curious that the simplest interpretation of the data that the filament itself was a very extended version of already known redox chains,^38^ but composed of OmcS, was considered less plausible than a cytochrome decorated pilus with hitherto unknown metallic-like electrical properties^10^ that required a “new paradigm”^10^ for biological electron transfer.

According to Malvankar and co-workers, the AFM and antibody results supported the identification of the pilus from the following pieces of genetic data:^3^ (1) The level of RNA for the pilus was high in electron transferring cells; (2) The genomic organization of the pilus biosynthesis genes was similar to what was found in other type IV pili-producing bacteria; (3) The primary sequence of the pilus resembled the N-terminal sequence from other type IV pili-producing bacteria; (4) A pili-deletion mutant strain lacked filaments and the ability to transfer electrons extracellularly; and (5) Heterologous expression of pili from other *Geobacter spp.* or point mutations in the *G. sulfurreducens* pilus caused cells to produce filaments that had a ∼10^7^-fold range in conductivity (3.8 × 10^-5^ to 2.8 × 10^2^S/cm) that correlated with aromatic residue density in the pilus.^18-20, 22^

The first three points are practically *non sequiturs* for identifying a protein, and the fourth point regarding pili deletion had the complicating factor of possible pleotropic effects. That explanation, however, cannot dismiss the conductivity-versus-aromatic density correlation in the fifth point that was discovered by Malvankar and co-workers.^22^ Out of a barrage of individually indirect or weak reasons for believing the filament was a pilus, this seems to be the only one that has not (yet) crumbled under the weight of an over-extended interpretation.

If Malvankar and co-workers’ new contention is right that filaments are cytochrome-based,^3^ then two important suppositions must be made: Genetic introduction of 5 different pili caused *G. sulfurreducens* to offer up 5 different cytochrome filaments, almost all with the same ∼3 nm diameter,^18-20, 22^ and (2) the conductivities of these pleiotropically expressed cytochrome filaments differed in precisely the way anticipated by the MLP hypothesis. Two of the putative pili for the correlation are now argued to be different cytochrome filaments,^3, 29^ and the true identities of the other filaments remain unknown.

The conductivity-versus-aromatic density correlation is presently unexplained in light of the CryoEM-resolved structure of the *G. sulfurreducens* pilus that shows no continuously stacked chain of aromatics for conductivity,^28^ as well as multiple cytochrome filament structures.^3, 23-27^ The present article demonstrates that the ∼10^7^-fold variation in conductivity is both physically implausible for a cytochrome architecture and physiologically irrelevant.

### 1.3. Is the filament Dumbo?

An array of characterizations substituted for sight before the CryoEM structure of any cytochrome filament was resolved. Those characterizations showed (1) a sigmoidal voltage dependence of the conductivity with greater conductivity at highly oxidizing potentials;^9, 13^ (2) crystallinity indicative^10^ of π-orbital overlap and long-range electron delocalization that linearly correlated with the aromatic density of the putative pilus and increased upon acidification;^17^ (3) carbon nanotube-like rapid propagation of holes over the entire structure that was independent of temperature, enhanced with acidification, and prevented by mutating out aromatic residues in the *PilA-N* gene;^15^ (4) a cooling-induced exponential increase, followed by an exponential decrease in conductivity,^9, 30^ and (5) a ∼10^3^-fold enhancement in conductivity upon proton doping down to pH 2^13^ that could not be explained if cytochromes were involved: “The pH dependence shows that pili behave similar to organic metals; there is no comparable pH-dependent model for cytochrome-based electron transfer.”^10^

All of these dependencies were interpreted as incompatible with the RCTC hypothesis, and to confirm the MLP hypothesis. If the structure of the filament was “everybody’s elephant in the room,” and now we know that elephant to be a cytochrome, one wonders if the elephant is of the flying variety (Dumbo): Do the cytochrome filaments have hallmarks of metallic-like conductivity hitherto unknown for other cytochromes that we need to learn to appreciate? Or are the characterizations simply erroneous? Is there any metallic-like conducting protein from *G. sulfurreducens* whatsoever? Figuring out whether any of the reported conductivity dependencies are relevant to the CryoEM structures of cytochrome filaments is of utmost urgency to spare the field another decade of trying to rationalize a pleasant vision for bioelectronics that may simply be a mirage.

The present article, therefore, asks: Are electrical characterizations of *Geobacter* ‘nanowires’ consistent with the CryoEM-resolved cytochrome filament structures,^3, 23-27^ theoretical work based on them,^30, 71-76^ electron transfer kinetics measured in other multi-heme proteins,^77, 78^ and biological considerations?^31, 79, 80^ The answer is shown to be “No” in almost every case, at least from the perspective of *in vivo* conditions. Instead, functionally robust, and physiologically relevant redox conduction is found to describe the mechanism of cytochrome filaments. It is suggested that the hallmarks of metallic-like conductivity are an artifact of the solid-state conditions of the experiments under which electron transport (not biological electron transfer) occurs, the structures resolved by CryoEM are not the same structures electrically characterized, or the electrical measurements to date are not trustworthy.

## 2. Methods

The present study principally re-analyzes previously published energetic parameters and kinetic rate constants computed for cytochrome filaments.^30, 71, 74^ These analyses were performed using the BioDC program^81^ developed by the present author, which is publicly available at the GitHub Repository: www.github.com/Mag14011/BioDC. BioDC includes an implementation of the analytical Derrida formula for single-particle diffusion along a one-dimensional periodic chain of hopping sites that was generously contributed by Fredrik Jansson.^82, 83^ BioDC also includes an implementation of the multi-particle steady-state flux kinetic model developed and kindly provided by Blumberger and co-workers.^71, 84, 85^

In Section 3.1.1.3, Constant Redox and pH Molecular Dynamics (C(E,pH)MD) simulations are reported in Figure 2. These simulations were conducted are previously described,^73^ except that the hemes in the flanking subunits of the trimer model were either held in the oxidized or reduced state while the hemes in the central subunit were titrated as a function of solution potential. These sets of simulations corresponded to cathodic and anodic electrochemical sweeps, respectively.

**Figure 2.**
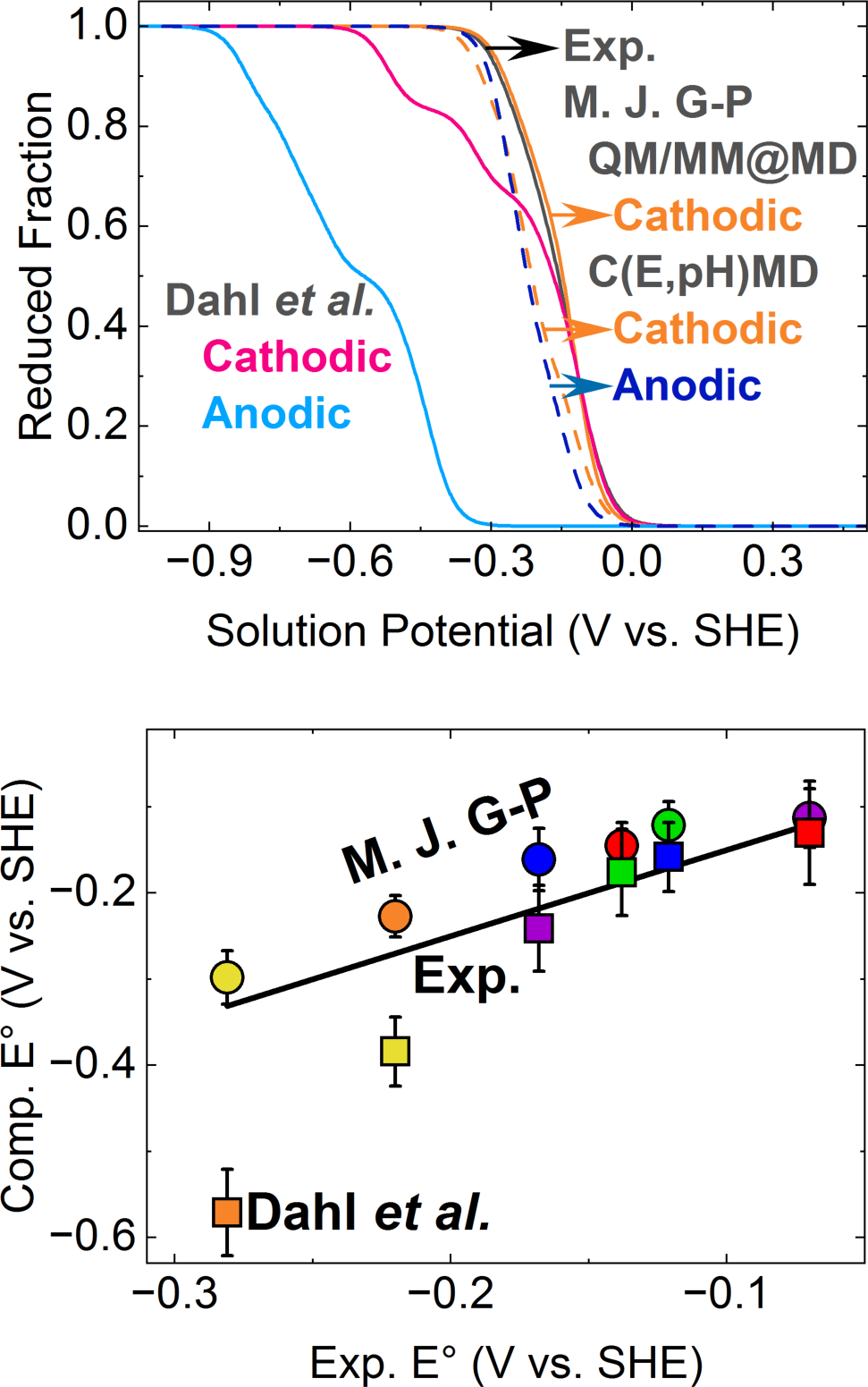
(*top*) Comparison of spectroelectrochemical^101^ and computed titration curves by Dahl *et al.^30^* and the present author (M. J. G-P)^73^ using either quantum mechanical/molecular mechanical computations at classical molecular dynamics-generated configurations (QM/MM@MD) or constant redox and pH molecular dynamics (C(E,pH)MD) computations. (*bottom*) Comparison of experimental macroscopic and simulated microscopic redox potentials (under cathodic conditions) presented as a correlation plot where deviation from the diagonal indicate divergence from the spectroelectrochemical-derived values.

In Section 3.2.2, powder XRD patterns were simulated using the freely available Mercury program^86^ from the Cambridge Crystallographic Data Center (CCDC).

## 3. Results and Discussion

### 3.1. Measured Conductivities are either Physiologically Irrelevant or Physically Implausible for Cytochrome Filament Structures

In this sub-section, physical constraints on the energetics of heme-to-heme electron transfer are established (Section 3.1.1). The associated kinetic rate constants are found to be largely independent of protein identity for highly conserved heme packing geometries (Section 3.1.2). The observation of conductivity in fully oxidized or reduced cytochrome filaments (Section 3.1.3), as well as the magnitudes of the reported conductivities (Section 3.1.4–3.1.5) cannot be accounted for by physiologically relevant redox conduction. Some of the discrepancies are attributed to physical artifacts of the experimental conditions including electrode adsorption, dehydration, compression-induced deformation, and the number of heme sites coupled to the electrodes (Section 3.1.5). The effective electron transfer rate constant for OmcZ is nearly at the non-adiabatic ‘speed limit’ if one heme site is assumed to be coupled at each protein-electrode interface, which is physically nonsensical for weakly coupled metal centers separated by 9–11 Å.

#### 3.1.1. Physical Constraints on Electron Transfer Energetics

Reduction-oxidation (redox) conduction is thought to describe the flow of electrons in multi-heme proteins under physiologically relevant conditions.^87, 88^ The kinetics of each successive redox reaction is generally well-described by non-adiabatic Marcus theory,^89, 90^ which defines the rate constant for each electron transfer in terms of three energetic parameters (Eq. 1): (1) The electronic coupling between the charge donor and acceptor (⟨H⟩), (2) the reorganization free energy (λ), and (3) the reaction free energy (ΔG^∘^).

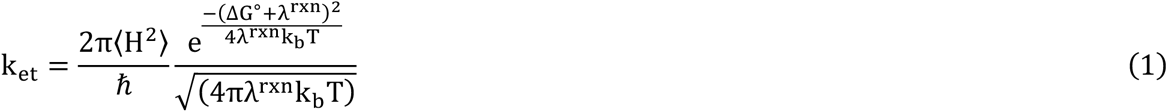

For a chain with N steps, there are 3N energetic parameters that need to be defined. For an electron to cross a 300-nm gap between electrodes via the cytochrome filament ‘bridge’, it must take ∼400 such heme-to-heme steps. Even if the homopolymeric nature of the Omc- S and Z filament is considered, 6 or 7 steps are needed to move between identical hemes in adjacent subunits along the main cofactor chain. Computations of the electron flux through the ‘unit cell’ of these homopolymers requires 18–24 parameters. Given the reference to elephants above, the large number of parameters brings to mind the problem of Von Neumann’s elephant: There are so many parameters that anything can be described. Are there any physical constraints, either expected or measured, that can be applied to these parameters?

##### 3.1.1.1. Electronic Coupling

The electronic couplings are principally determined by the distance and orientation (or packing geometry) of the heme cofactors.^91, 92^ The surrounding environment has a perturbative (∼10%) influence.^73, 92^ Heme packing geometries in all known cytochromes are highly conserved among multi-heme proteins.^24, 25^ The topologies come primarily in two different flavors: In the parallel-displaced (“slip-stacked”) configuration, adjacent heme macrocycles are nearly parallel (tilt angle < 20°), separated by ∼3–5 Å (edge-to-edge minimum distance), and rotated by ∼180° with respect to each other. In the perpendicular (“T-stacked”) configuration, adjacent heme macrocycles are nearly perpendicular (tilt angle ≈ 90°), separated by ∼4–6 Å, and rotated by ∼110–140° with respect to each other.

The heme-packing parameters apply to almost all heme pairs in cytochrome filaments and they are well maintained over hundreds of nanoseconds in classical molecular dynamics simulations at 300 K (Table S1).^74^ Only two heme pairs in OmcZ deviate from these classifications: The heme that branches off from the main chain has a rotation angle of 56° with respect to the nearest main chain heme, and that heme is rotated by 82° with respect to a neighboring heme in the main chain.^25^ These heme pairs have edge-to-edge separations that are ∼1.0 Å smaller than expected from a survey of >800 cytochromes in the PDB.^24, 25^

Still, heme-to-heme electronic couplings in multi-hemes examined to date,^85, 93^ including the cytochrome filaments,^30, 71, 73, 74^ and regardless of theoretical method have electronic couplings ≤0.016 eV (Table S2). The tunneling pathways between hemes may be through empty space, protein backbone bonds, or inter-heme (His axial ligand-to-propionic acid side chain) H-bonds,^74^ but the variation in couplings is still within a factor of 5.^74^ Weak (sub-thermal energy at 298 K) electronic couplings between T- and slip-stacked heme pairs are independent of multi-heme protein identity.

Regarding OmcZ specifically, the coupling for T- and slip-stacked heme pairs were shown to be 0.7 and 1.8 times the respective values in OmcS.^74^ Because the changes from Omc- S to Z are relatively small and of a mixed (increased/decreased) nature, there is no compelling support for Malvankar and co-workers’ (including the present author)^26^ hypothesis that larger inter-heme electronic couplings explain the 10^3^-fold greater conductivity of OmcZ. This result was omitted from that publication, which proports in the title to identify the mechanism of conductivity. The only coupling that is likely enhanced in OmcZ is the protein-electron coupling via an almost entirely solvent exposed heme.^25, 26^

##### 3.1.1.2. Reorganization Energy

The reorganization energies for heme-to-heme electron transfers are expected from previous studies^88^ and computed specifically for the cytochrome filaments^30, 71, 73, 74^ to be ∼0.4–1.0 eV (Table S3). These reorganization energies may be overestimated by ∼20–40% because polarizability of the environment was not included in classical molecular dynamics simulations.^94,95^ Polarizability of the heme active site may further lower the reorganization energy, as proposed for cytochrome *c* on the basis of semi-empirical quantum chemistry (ZINDO/S) calculations,^96^ although this point has been debated.^97-99^ However, density functional theory calculations for the hemes in the cytochrome filaments found only <0.04 eV shifts in the reorganization energy when relaxing vacuum-optimized electron densities in the presence of the perturbing protein/water environment (Tables S4–S6.^74^

Polarizability corrections are not expected to vary widely between proteins, as evidenced by recommendations of system-independent scaling factors.^94, 95^ These corrections may shift the conserved range of reorganization energies from ∼0.4–1.0 to ∼0.3–0.8 eV or lower. Regardless, however, the reorganization energies remain of the same order of magnitude (tenths of an eV). Besides providing a protein-independent expectation for computed reorganization energies, this magnitude is at least 10-fold greater than the expected electronic couplings, indicating that non-adiabatic Marcus theory is the appropriate description for heme-to-heme electron transfer in multi-heme proteins.^87^

##### 3.1.1.3. Reaction Free Energy

Reaction free energies reflect the difference in redox potential between the charge donor and acceptor. The redox potentials for bis-histidine ligated *c*-type hemes are expected from a wealth of experimental studies to fall in the range -0.350 to +0.150 V versus standard hydrogen electrode (SHE).^100^ Thus, the maximum expected free energy difference between two bis-histidine-ligated *c*-type hemes is |0.5| eV. However, it is unlikely that second-sphere interactions can create a 0.5 V difference between two adjacent hemes in van der Waals contact. The computed free energy differences for cytochrome filaments do not exceed |0.26| eV^74^ (except for the work by Dahl *et al.*;^30^ see below for an explanation) and are usually ≤||0.2| eV (Table S7).^71, 74^ These values are consistent with the expectation that reaction free energies are kept small (close to zero) in biology. The physical constraints on simulated free energies can be made tighter in the case of OmcS. A spectroelectrochemical analysis^101^ of OmcS filaments recently became publicly available. The analysis shows that the experimental reduced fraction-versus-solution potential titration curve can be approximated by a sum of six independent or coupled) Nernstian equations, one for each heme in the subunit of the homopolymer.^101, 102^ A set of potentials for the multi-redox center protein (macroscopic potentials) were obtained that can be related to the cofactor-specific (microscopic potentials) if inter-cofactor interactions are sufficiently small. As discussed in the *Note on Interpreting Spectroelectrochemical Titration Curves* in the Supporting Information, heme-heme interactions are expected from experiments on other multi-heme proteins^103^ and synthetic diheme model systems,^104^ as well as computations on OmcS^73^ to be comparable in magnitude to the sub-0.1 V uncertainties in the computed redox potentials.

Figure 2 shows the experimental and simulated titration curves, as well as the comparison of experimental macroscopic and computed microscopic potentials. The redox midpoint for the OmcS filament is predicted within 0.010 V and the experimental macroscopic and computed microscopic potentials agree within ±0.038 V (Table S8), suggesting that one particular heme dominates each redox transition in the filament. This level of agreement on six independently predicted redox potentials from quantum mechanical/molecular mechanical computations at classical molecular dynamics-generated configurations (QM/MM@MD) is remarkable.^105^

The only other set of available heme redox potentials computed for OmcS come from Dahl *et al.* (including the present author who advised on that work).^30^ As shown in Figure 2 (*top*), there are large discrepancies between the experimental titration curve, which does not show an appreciable hysteresis, and those simulated for cathodic and anodic electrochemical sweeps using the redox potentials reported by Dahl *et al*. (Table S8). The experimentally false difference in the computed redox profile under cathodic and anodic conditions, which was known up to two years *before* publication,^101, 106^ was used to justify why dithionite-reduced versus air-oxidized OmcS is more conductive.^30^

Figure 2 (*bottom*) and Table S8 show that two of the redox potentials reported at 310 K under cathodic conditions by Dahl *et al.* deviate from experiment by 0.11 and 0.24 V, which, again, was known to be the case up to two years *before* publication but never disclosed to reviewers/readers.^101, 106^ As discussed before^73^ (and summarized in Section 3.2.4 below), the physically unrealistic and experimentally false redox potentials allowed Dahl *et al.* to severely underestimate the redox conductivity at high temperature, and thereby *seemingly* validate the experimental observation of greater conductivity at lower temperature; the “right” result for a fictitious reason. The Dahl *et al.* study has neither been retracted nor corrected, or followed-up by a subsequent work to correct the record except independently by the present author.^73^

##### 3.1.1.4. Summary

The energetic parameters for heme-to-heme electron transfer, to first approximation, are generic to the class of multi-heme proteins: Typically, couplings are ≤0.02 eV; reorganization energies are ∼0.4-1.0 eV, and reaction free energies are <|0.5| eV. This view is actually the standard perspective for electron transfer in proteins, but sometimes needs to be corrected for polarizability^96^ and/or dynamical effects.^107^

All these physical constraints are satisfied by computations published to date (Tables S2, S3, S7). For example, the present author found previously that the electronic couplings, reaction free energies, and reorganization free energies for the Omc- E, S, and Z filaments were respectively <0.015 eV, <|0.28| eV, and 0.48 – 0.98 eV.^74^ As concluded in that study, the redox-based conduction of electrons along the heme chains in cytochrome filaments is functionally robust to structural diversity. It should be expected, then, that the rates computed from these physically reasonable energetic parameters agree well with analyses of electron transfer kinetics in multi-heme proteins. This result is precisely what is demonstrated in the next section.

#### 3.1.2. Heme-to-Heme Electron Transfer Rates are largely Independent of Protein Identity

Kinetic analyses of ultrafast transient absorption measurements on photosensitized variants of the metal reducing cytochrome type C (MtrC)^78^ and the small tetraheme cytochrome (STC)^77^ from *Shewanella oneidensis* have provided experimental estimates for heme-to-heme electron transfer rates between parallel-displaced (slip-stacked) and perpendicular (T-stacked) heme pairs. The average forward/backward rates were 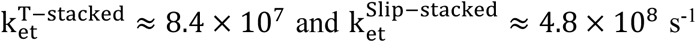. The slowest and fastest rates for the T-stacked geometry (8.7 × 10^>^ − 1.3 × 10= s^-1^) were within a factor of 10 slower than the slowest and fastest rates for the slip-stacked geometry (1.4 × 10^8^ − 1.6 × 10^9^ s^-1^).

Blumberger and co-workers proposed that these ∼10^7^ – 10^9^ s^-1^ heme-to-heme electron transfer rates are likely generalizable to other proteins that have the same highly conserved slip- and T-stacked packing geometries.^78^ Two decades earlier, Page, Moser, and Dutton proposed that the proximity of redox centers alone was sufficient to ensure electron transfer rates were largely insensitive to the surrounding protein and faster than typical enzymatic turnover.^108^ Figure 3 (Tables S9) evaluates these suggestions by comparing the experimental rates from MtrC and STC to previously computed rates for Omc- E, S, and Z from the present author,^74^ OmcS from Jiang *et al.*,^71^ and OmcS from Dahl *et al.*,^30^ as well as the typical range for enzymatic rates.^80^

**Figure 3.**
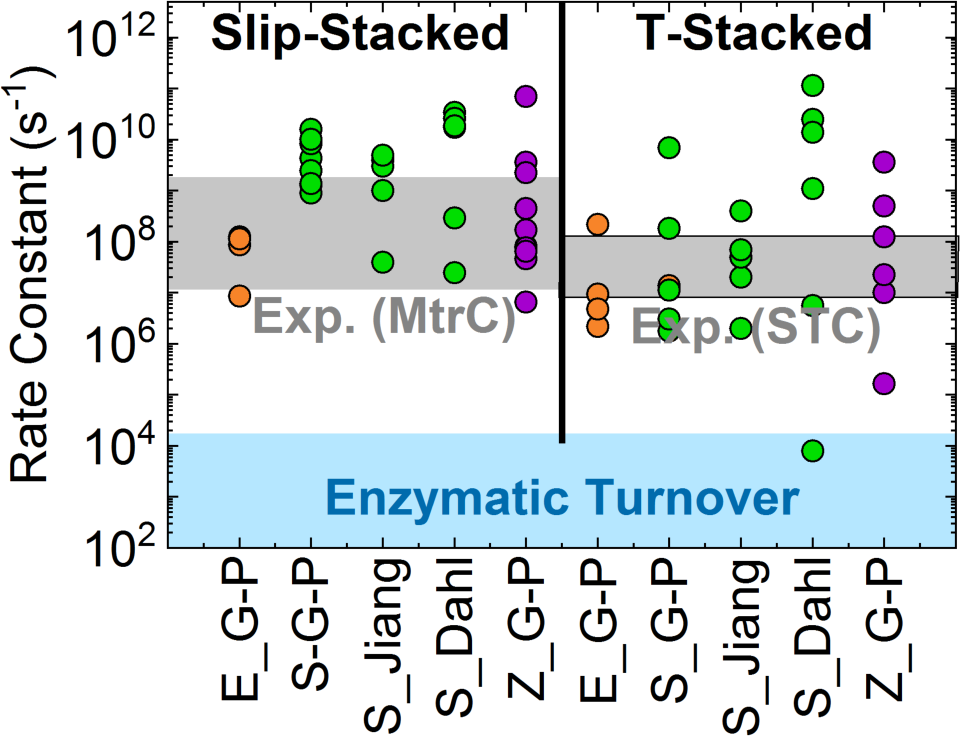
Comparison of heme-to-heme electron transfer rates previously measured in MtrC^78^ and STC^77^ by Blumberger and co-workers to rates computed for Omc- E, S, and Z by the present author,^74^ OmcS by Jiang *et al*.^71^ and OmcS by Dahl *et al.*,^30^ as well as the typical range for enzymatic turnover.^80^ The common “Omc” part of the protein names is omitted in labels, and the present author’s name (Guberman-Pfeffer) is abbreviated as G-P for conciseness.

The slowest, fastest, and average computed rates in Omc- E, S, and Z for either the slip- or T-stacked geometries come within a factor of <10^2^ of the same metrics for the experimental rates in MtrC and STC (Table S10). More specifically, the slowest, fastest, and average computed rates for T-stacked heme pairs in all three filaments come within factors of 22, 67, and 23, respectively, of the same metrics for STC;^77^ the slowest, fastest, and average computed rates for slip-stacked heme pairs in all three filaments come within factors of 91, 9, and 6, respectively, of the same metrics in MtrC.^78^ These observations apply to the works of Jiang *et al.^71^* and the present author,^74^ but not to Dahl *et al.*,^30^ which reported slowest, fastest, and average rates for T-stacked heme pairs that deviate by 3000, 554, and 282-fold, respectively, from the same metrics reported experimentally for STC.^77^ The problem is related to much too large free energy differences because of unphysically negative and experimentally false^101, 106^ computed redox potentials (Figure 2), as discussed above.

The level of agreement for all theoretical work except by Dahl *et al.* is strikingly of a similar order of magnitude as previously found when directly comparing computed and measured rates for MtrC^78^ and STC.^77^ Note that those computations did not include the presence of the Ru-dye; the agreement with experiment therefore implies that the dye did not significantly alter the intrinsic electron transfer rates. Thus, as anticipated, heme-to-heme electron transfer rates within highly conserved packing geometries are largely transferrable among multi-heme proteins, and experiment and theory are in excellent agreement on the magnitude of the rates.

Another important conclusion from Figure 3 is that the rate-limiting step in a chain of both slip- and T-stacked heme pairs, as found in all known cytochrome filaments, is likely to be ∼10^5^ – 10^7^ s^-1^ and belong to a T-stacked heme pair. This conclusion extends to A3MW92 and F2KMU8, filaments which were analyzed recently with BioDC,^81^ a highly automated workflow for computing redox conductivity implemented by the present author. BioDC uses the heme packing geometry, solvent accessibility, and the difference in electrostatic energies for the reduced and oxidized states from Poisson-Boltzmann calculations to estimate the energetic parameters for Marcus theory electron transfer rates, which in turn are used to compute diffusive single-particle and steady-state multi-particle fluxes.

That all cytochrome filaments have half^3, 23, 24, 27^ or nearly half^25, 26^ of the heme pairs in a geometry that is a bottleneck for electron flow indicates that these structures are not optimized by evolution to carry the greatest amount of current. As suggested previously^74^ and shown in Figure 3, intra-filament electron transfer is already faster than typical enzymatic turnovers (≥ µs vs. ms). The implication is that the rate-limiting step for respiration resides elsewhere in the cellular machinery and there is no biological advantage for further optimizing the electrical conductivity of the filaments.

There, however, is evolutionary pressure to adapt the environment-filament interface formed by the protein sheath insulating the heme chain; for example, to endure mechanical stresses and to bind different partners for exchanging electrons. From this perspective, the filaments comprise a functionally robust heme chain wrapped in environment-customized protein packaging. The ‘design’-strategy was previously analogized to how the conserved photosystems of photosynthesis are interfaced to distinct spectral niches by highly adapted light harvesting antenna.^74^

To summarize, heme-to-heme electron transfer rates within highly conserved packing geometries are also highly conserved, typically falling within the 1 – 10 ns timescale as proposed by Blumberger and co-workers.^78^ In this view, the observed ∼10^7^-fold variation in the conductivity of filaments obtained by introducing genetic pili variants into *G. sulfurreducens* cannot be explained if, as Malvankar and co-workers suggest,^3^ those filaments are now to be interpreted as cytochromes. The variation does not seem biologically relevant considering Figure 3.

The generic rates are used below to assess the limits of redox-based conductivity in cytochrome filaments and whether or not the reported conductivities are physically plausible. Before doing that, however, the question of whether the non-zero conductivities of the filaments reported in fully oxidized or fully reduced redox states is consistent—regardless of its magnitude—with the physiological process of redox conduction.

#### 3.1.3. Conductivity in Fully Oxidized or Reduced States Inconsistent with Redox Conduction

Malvankar and co-workers previously reported a sigmoidal voltage dependence of the conductivity for the wild-type *G. sulfurreducens* pilus (GsP), with the conductivity increasing by >10^2^-fold at highly oxidizing potentials.^13^ The observation *seems* consistent with the theoretical prediction that strongly oxidizing conditions are needed for pili conductivity.^55^ These observations were inconsistent with redox conduction through cytochromes. Redox conductivity is expected to be sharply peaked at the formal redox potential and rapidly fall to zero at the extremes of fully oxidized or fully reduced, because no heme would be available to donate or accept electrons, respectively.^7^

More recently, Malvankar and co-workers reported that fully air-oxidized Omc- S and Z (said to be previously misidentified as GsP^3^ and its F51W-F57W mutant (GsW51W57)^29^, respectively) ‘nanowires’ have conductivities of ∼0.05 S/cm^3, 18^ and ∼4–30 S/cm^19, 29^ at pH 7. OmcS filaments were additionally reported to be 6-fold more conductive in the dithionite-reduced vs. air-oxidized state.^30^ These observations are *still* inconsistent with redox conduction through cytochromes for precisely the same reason.

A reduced (oxidized) population of hemes is not expected to form under the applied bias in the experiments—and there is no direct evidence that suggests otherwise—because redox reactions require ion mobility in a surrounding electrolyte,^7, 109^ but the experiments were performed on air-dried samples.^29, 30^ UV-vis spectra collected for a film of fully oxidized OmcS filaments before and during application of an external bias showed only a small (≤5 nm) shift—primarily in the tails of a >60 nm wide peak—towards the spectral feature indicative of a reduced population of hemes.^30^ The shift may reflect voltage-induced changes in the energy levels probed by photoexcitation instead of formal reduction of the hemes by the applied bias. The relatively small spectroscopic shift suggests that abiological electron transport (ETp) is more likely than physiologically relevant electron transfer (ET) under the experimental conditions of bear Au electrode-adsorbed and air-dried filaments, because solid-state ETp does not (whereas biological ET does) depend on the redox activity of the heme-Fe center,^87, 110-112^ and the presence/absence of water can significantly change the conductivity.^113^

Dahl *et al.^30^* attempted to explain the increased conductivity of chemically-reduced OmcS with computations that showed—in contradiction to spectroelectrochemical data from the same laboratory (Figure 2)^101, 106^—a 0.3–0.6 V hysteresis in the redox potential for each heme. Anantram and co-workers proposed^76^ instead that the quantum transmission function is increased in the reduced state at lower Fermi energies because of the redox-linked shift in orbital energies. The alignment of the orbitals with the Fermi level of the electrodes will change upon reduction, and greater protein-electrode coupling is possible.^114-117^ If true, this effect is then not relevant to biology.

With regard to the sigmoidal voltage dependence observed for what were thought to be GsP and now claimed to be OmcS, either the new attribution to a cytochrome is wrong, or the experiments were performed incorrectly, as previously argued^10, 21, 44, 57^ because the observation seems irreconcilable with the redox-active nature of a cytochrome.

#### 3.1.4. Limits of Physiologically Relevant Redox Conductivity

Using 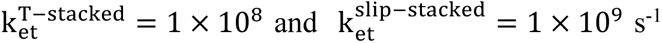 as order-of-magnitude estimates from Section 3.1.2., Figure 4 compares the predicted steady-state current as a function of filament length for heme chains that are (1) purely slip-stacked, (2) purely T-stacked, (3) strictly alternating slipped and T-stacked as found in OmcS,^3, 23^ OmcE,^24^ A3MW92,^27^ and F2KMU8,^27^ or (4) follow the T → S → S → T → S → T→ S (S = slipped) stacked pattern along the main chain of OmcZ (referred to as “mixed”).^25, 26^ Figure 4 compares these theoretical limits to currents predicted by structure-based calculations using BioDC^81^ and QM/MM@MD techniques,^74^ as well as the experimentally reported^3, 29, 30^ currents for cytochrome filaments.

**Figure 4.**
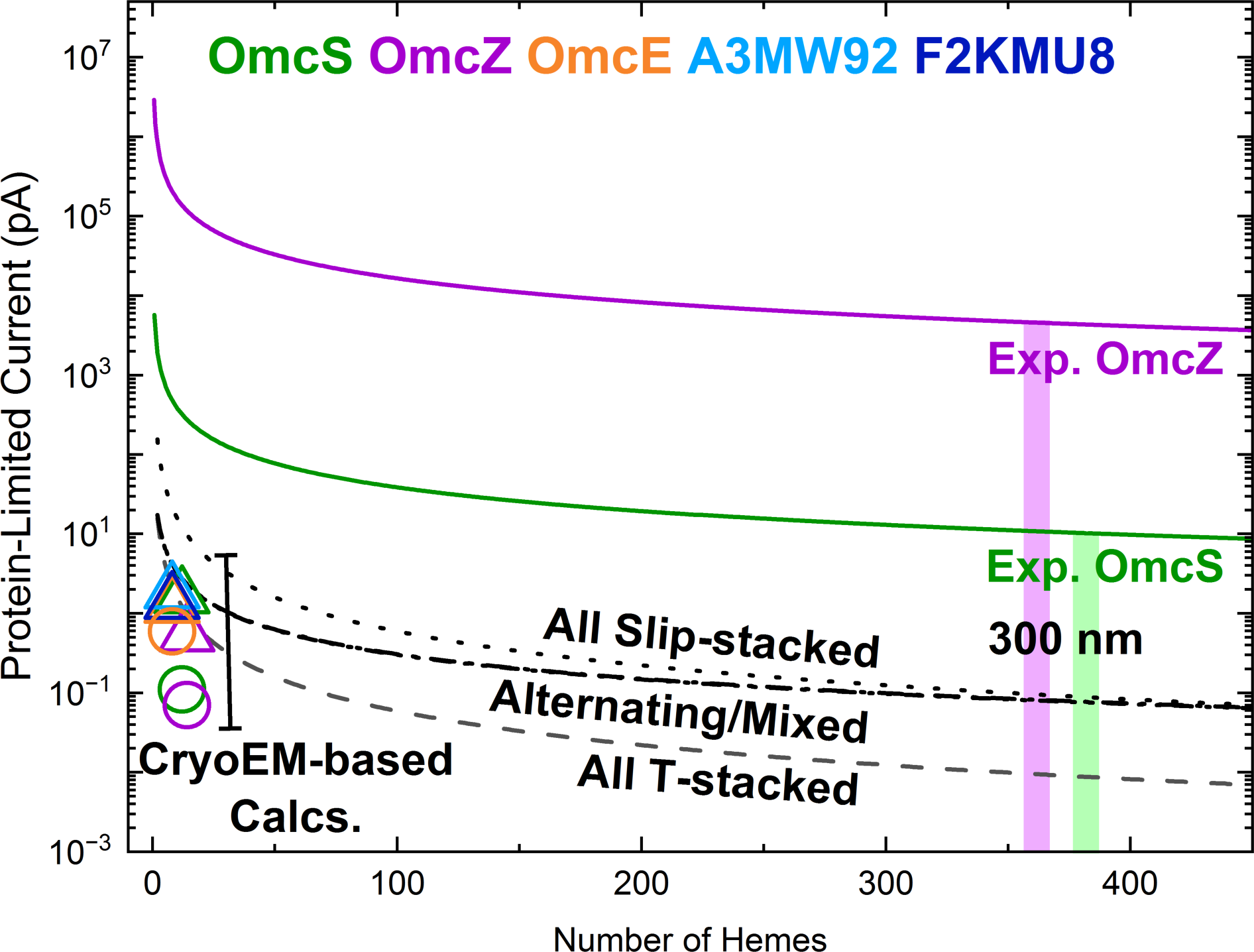
Comparison of experimental and theoretical electrical currents in cytochrome filaments. The experimental currents are computed at an applied bias of 0.1 V using the reported conductivities and filament dimensions.^3, 29, 30^ The computed currents for all known cytochrome filaments using BioDC (triangle) and/or QM/MM@MD (circle) techniques are based on previously reported Marcus electron transfer rates.^74, 81^ The computed currents for hypothetical All Slip-stacked (dotted line), All T-stacked (dashed-line), and Alternating or Mixed T/Slip-stacked (dash-dot and dash-dot-dot lines) use generic order-of-magnitude estimates for the electron transfer rate constants of k^slip-stacked^ = 1 × 10^9^ and k^T-stacked^ = 1 × 10^8^ s^-1^. Note that the curves for the Alternating and Mixed cases virtually superimpose. The curves for the hypothetical cases are fits to computed currents for chains with 18 or fewer hemes according to the equations 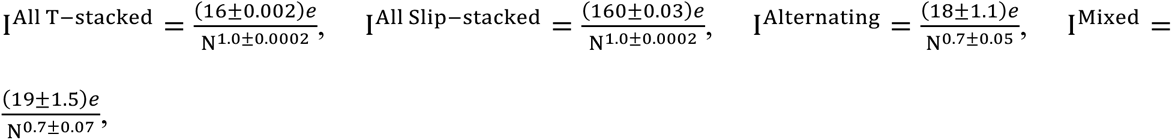 and correlation coefficients of 1.0000, 1.0000, 0.9264, and 0.8595, respectively.

The computed currents in Figure 4 were obtained as protein-limited (*i.e.*, electron injection/ejection rates exceeded inter-heme rates) steady-state electron fluxes (J) using the kinetics model of Blumberger and co-workers.^71, 84, 85^ The fluxes were converted to currents by the first term on the right-hand-side of Eq. 2. This approach is increasingly harder to apply as system size increases because of time demands and convergence issues with iteratively solving coupled polynomial equations. Systems with 18 or fewer hemes (N_hem_ ≤ 18) were therefore directly computed (Table S11) and fitted to the expected distance dependence for hopping transport (second term on the right-hand-side of Eq. 2).^118^

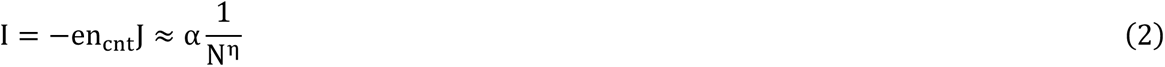

where e is the electron charge and n_cnt_ is the fewest number of contacts at either protein-electrode interface,^56^ momentarily assumed to be 1.0; that n_cnt_must be greater than this value, as shown below, is an important result of the present study. As for the fitted curves, α is a proportionality constant, N is the number of charge-relaying heme-to-heme steps, and η can take values from 1 (irreversible) to 2 (reversable) hopping steps.^119^ The fitted curves for the All Slip-stacked, All T-stacked, Alternating T/slip-stacked, and Mixed (OmcZ-style) T/slip-stacked cases in Figure 4 had correlation coefficients of 1.0000, 1.0000, 0.9264, and 0.8595, respectively.

The experimental currents in Figure 4 were computed from the reported conductivities and the CryoEM geometries of the filaments by Eq. 3, which is essentially Ohm’s law.

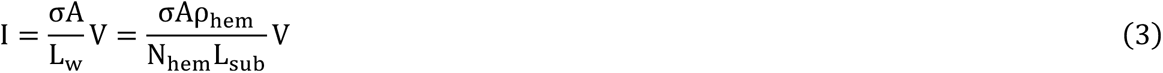

where σ is the conductivity (0.024 and 27.8 S/cm for Omc- S and Z, respectively, at pH 7), ^29, 30^ A is the cross-sectional area of the conduction channel, which is approximated as πr^#^ for a cylinder with r_c_ taken to be the radius of the heme group (7.5 × 10^-8^ cm).^72^ Note that experimental reports^3, 29, 30^ have instead used the half-height of the filament from atomic force microscopy experiments (2.0 × 10^-7^and 1.3 × 10^-7^ cm),^29, 30^ but this choice assumes that the entire filament (protein+heme) is the conduction channel and that the AFM measurement is not rendered meaningless because of dehydration artifacts.^120^ The present author has demonstrated some contribution of the protein to tunneling pathways between the hemes, particularly in OmcZ,^74^ but not enough to justify assuming that the entire protein participates in the electrical conduction under physiological conditions, which is the concern here. Molecular orbitals distributed over the protein, however, can play an important role in abiological ETp.^112^

L_W_ is the length of the filament, which is expressed as the total number of hemes (N_hem_) divided by the number of hemes-per-subunit along the main chain (ρ_hem_; 6 for OmcS and 7 for OmcZ) and multiplied by the length of a subunit (L_sub_; 4.7 × 10^-7^ and 5.8 × 10^-7^ cm for Omc- S and Z, respectively). Note that the heme that branches off from the main chain in OmcZ was not counted because it only siphons electrons into a ‘dead-end’ so that soluble species can be reduced. By neglecting the branched nature of the heme chain in OmcZ, the simulated current along the main chain will be an over-estimate, which makes the under-estimation that is found with respect to experiment (see below) even larger.

V is the applied bias, taken to be a physiologically relevant 0.1 V. However, it should be noted that the potential difference is not experienced in a physiologically relevant way in the electrical measurements. If the potential difference between the half reactions inside and outside the cell falls linearly across the micron-long filament under physiological conditions (biological ET regime), the potential drop is less than thermal energy between any pair of hemes, and it is also screened by mobile ions.^7, 56^ The electrons move through the filament according to concentration gradients of reduced and oxidized species, not the electrical potential difference between the half reactions.^7, 109^ Under the solid-state conditions of the electrical measurements (abiological ETp regime), however, the seemingly physiologically relevant bias is not effectively screened by ions with limited or no mobility. The applied bias alters the free energy landscape from what it would be under physiological conditions and electrons move according to electrical, not chemical potential differences.^7, 109^

At a potential of 0.1 V, the *Note on Intrinsic versus Contact Resistance* in the Supporting information establishes that the experimental currents are limited by the resistance of the protein instead of the protein-electrode contacts when the filaments are >30 nm long, or equivalently contain >36 hemes along the main chain.

For a filament that spans a 300-nm gap between electrodes, as used experimentally, the CryoEM structures of Omc- S and Z suggest there are ∼400 hemes. At this length, the currents predicted from the experimental rates in MtrC and STC range from ∼10^-2^ pA for purely T-stacked to ∼10^-1^ pA for purely slip-stacked. The predicted current through an alternating or OmcZ-style mixed T/slip-stacked chain, interestingly, is undistinguishable from the purely slip-stacked heme chain at long distance. The currents simulated for short (two subunit) filaments from the CryoEM structures using either BioDC or QM/MM@MD techniques for all known filaments fall in the vicinity of the mixed or purely T-stacked curves in Figure 4. The overall flux is limited by the T-stacked heme pairs. Thus, Figure 4 shows that the fluxes computed from the experimental heme-to-heme rates in MtrC and STC provide good approximations to the fluxes computed by applying theory to the CryoEM structures of the cytochrome filaments with the same heme packing geometries.

At odds with these experimental expectations and theoretical predictions are the ∼11 pA and ∼5 nA currents experimentally reported at an applied bias of 0.1 V for Omc- S and Z bridging a 300 nm gap, respectively.^3, 29, 30^ Contrary to some^30^ but not all^72-74^ prior claims, Figure 4 suggests that a physiologically relevant succession of redox reactions (multi-step hopping) cannot even come close to account for the reported conductivities of the cytochrome filaments. The next section shows that this conclusion applies generally to the conductivities measured for filaments from *G. sulfurreducens*, but the discrepancy may be ameliorated to some extent by considering physical artifacts of the experimental conditions.

#### 3.1.5. Unphysically Large Electron Transfer Rates may be an Experimental Artifact

Another way to assess the physical plausibility of the reported conductivities is by computing the effective hopping rate constant^55^ (k_eff_) and comparing to theoretical limits^56, 121-123^ and kinetic experiments for heme-to-heme electron transfer rates.^77, 78^ The analysis in this section is extended to consider the five filaments for which a ∼10^7^-fold change in conductivity was claimed to linearly correlate with aromatic residue density.^18-20, 22^ These filaments were assumed to be the wild-type *G. sulfurreducens* pilus with five aromatic residues (F24, Y27, Y32, F51, and Y57) mutated to Ala (GsAro5), the heterologously expressed *G. uraniireducens* pilus (GuP), the wild-type *G. sulfurreducens* pilus (GsP), the wild-type *G. sulfurreducens* pilus with Phe-51 and Tyr-57 mutated to Trp (GsW51W57), and the heterologously expressed *G. metallireducens* pilus (GmP). The GsP and GsW51W57 pili are now argued to be Omc- S and Z filaments, respectively.^3, 29^ For purposes of comparison, that argument is extended here to suppose that all these filaments were cytochromes of some sort. The reported conductivities were 4 × 10^-5”^ (GsAro5),^18^ 3 × 10^-4^ (GuP),^20^ 4.6 × 10^-2^ (GsP),^18^ 1.0 × 10^1^ (GsW51W57),^19^ and 3.0 × 10^2^ (GmP)^22^ S/cm.

It is straightforward (Eq. 4) to convert the conductivities to k_eff_ using (1) the definition of conductivity (σ) in terms of charge carrier density (ρ) and mobility (µ), (2) the Einstein-Smoluchowski expression relating µ to a diffusion coefficient (D), and (3) the definition of D in terms of a rate constant for a uniform chain with an average spacing of r_nm_between any two charge-relaying sites n and m. In Eq. 4, k_b_, T, and e are the Boltzmann constant, absolute temperature, and the magnitude of the electronic charge, respectively.

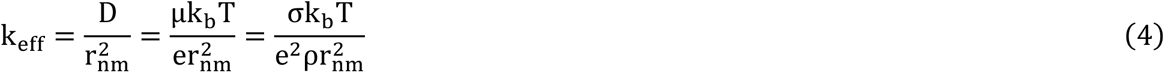

A lower-bound estimate for k_eff_ is obtained by assuming a maximal value for ρ. The maximal ρ that corresponds to redox conduction through a cytochrome occurs when every other heme is reduced 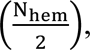 and the radius of the conduction channel (r_c_) is restricted to the radius of the heme group. In this case, and under the approximation that a cytochrome filament is cylindrical, 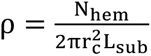, where N_hem_ and L_sub_ are the number of hemes per subunit and the length of a subunit, respectively. Setting r_C_to the radius of the heme group, the present author^74^ and others^72^ have estimated ρ to be ∼4 × 10^-20^ electrons/cm^3^ for Omc- E, S, or Z, which is of the same order of magnitude as previously measured for what were thought to be GsP.^15, 59^ As mentioned above, the choice of r_c_ seems sensible for a cytochrome because the hemes are the redox active charge-relaying sites in the filaments.

The focus here is on determining if the conductivities measured experimentally are physiologically relevant or even physically plausible.^112^ To that end, it is critical to realize that k_eff_ obtained with Eq. 4 is an *effective* rate constant because it is the rate at which charge transfers would need to occur in a uniform chain of sites (*i.e.*, all forward and backward rates are equal), whereas the measured or computed σs correspond to heterogeneous chains.

The cytochrome filaments have a mixture of T-stacked and slip-stacked heme pairs along the cofactor chain. The electron transfer rate between T-stacked hemes is, on average, a factor of 10 smaller than the rate between slip-stacked hemes (Table S10). The overall rate through a heterogeneous chain of multiple reactions in series is restricted to the rate of the slowest (rate-limiting) step. Thus, the effective rate constant for a homogeneous chain that gives the same conductivity as a heterogeneous chain reflects the rate-limiting step; individual steps in the heterogeneous chain may be faster. This is another reason—in addition to assuming a maximal value of ρ—why k_eff_ is a lower bound estimate for the electron transfer rates.

Applying Eq. 4 to the reported filament conductivities gives k_eff_s of 6 × 10^6^ (GsAro5), 5 × 10^7^(GuP), 8 × 10^9^ (GsP), 2 × 10^12^ (GsW51W57), and 5 × 10^13^ (GmP) s^-1^. These k_eff_s imply that the rate-limiting heme-to-heme electron transfer step—presumably for a T-stacked pair—is 10^-2^, 10^-1^, 10^1^, 10^4^, and 10^5^-fold faster than expected from kinetic analyses on other multi-heme proteins^77, 78^ or computed for structurally characterized cytochrome filaments.^71, 74^ Where CryoEM analyses are available, there is no structural basis for the unusually fast rates for GsP/OmcS and GsW51W57/OmcZ.

To consider the experimentally-derived rates from a more general perspective, the electron transfer rate can be described by a hopping mechanism if Eq. 5 is satisfied^122^ or band theory if Eq. 6 is satisfied.^56^

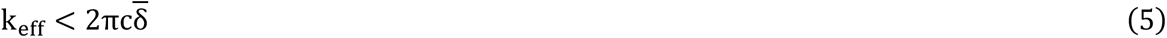

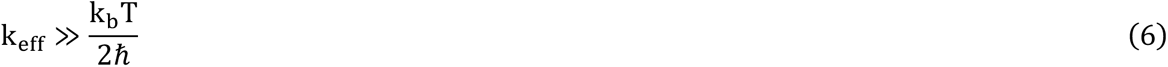

Where c is the speed of light and 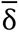 is the Raman line broadening as 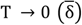, taken to be a ‘generous’ value of 3 cm^-1^ for molecular materials. Eqs. 5 and 6 indicate that k_eff_ must be less than 6 × 10^11^ s^-1^ for the hopping mechanism to apply, whereas k_eff_ must be much greater than 2 × 10^13^ s^-1^ to be described by band theory. Thus, electron transfers in the filaments known as GsAro5, GuP, and GsP can be described by electron hopping, whereas the electron transfers in GsW51W57 and GmP reside in the solvent-controlled dynamical regime between a hopping and band-theory description.^107^ Mobile electrons *appear* to propagate through GsW51W57 and GmP too quickly for thermal equilibration of the environment to make the hops independent (memoryless).^122^

Because GsP and GsW51W57 are respectively claimed^3, 29^ now to be the Omc- S and Z filaments for which the structures are known, the following subsections consider (1) If a hopping mechanism applies to electron transfers in OmcS, why have all theoretical works using a hopping mechanism based on the CryoEM structure^71-74^ failed to reproduce the experimentally-derived k_eff_? And (2) Is it physically plausible for the rate-limiting step in OmcZ to have a rate on the cusp of a band theory description?

##### 3.1.5.1. Number of Hemes Coupled to the Electrodes can Largely Rescue Hopping Models for OmcS

The experimentally derived k_eff_ is 8 × 10^9^ s^-1^ for OmcS. The k_eff_s found by Eq. 4 using theoretically computed charge diffusion coefficients are 6.4 × 10^6^ (Jiang *et al.*),^71^ 1.1 × 10^7^(present author),^74^ or 2.5 × 10^4^ s^-1^ (Dahl *et al.*).^30^ The result for Dahl *et al.* has the worst agreement with the experimentally-derived k_eff_ because of the erroneous free energy barrier discussed above. More interesting is the observation that the k_eff_s computed by Jiang *et al.*^71^ and the present author^74^using methods at very different levels of sophistication agree within a factor of <2. These k_eff_S underestimate the experimentally derived k_eff_by ∼10^3^-fold, but a physical artifact of the experimental conditions may account for a factor of ∼10^2^-fold.

Consider that the measured current (and thereby the conductivity and effective rate constant derived from it) is directly proportional to the fewest number of sites coupled to either electrode.^56^ Unlike partner redox proteins in biology, the electrodes cannot exchange electrons with the filament at the site of a specific heme. The “tip” used in the conducting probe atomic force microscopy experiments, for example, is 40–60 nm wide;^29, 30^ approximately 10 subunits of either Omc- S or Z can fit along its diameter. If only the heme sites are coupled to the electrodes, there are ∼10^2^ electron injection sites. The k_eff_s computed by Jiang *et al.* and the present author for OmcS assumed only one injection site. To be comparable to experiment, the predicted k_eff_s need to be scaled up by ∼10^2^-fold, leaving only a 10-fold underestimation.

The level of agreement realized for OmcS by accounting for the physical dimensions of the protein/electrode interface is pleasing, but probably fortuitous. The same argument regarding ∼10^2^ heme sites coupled to either electrode applies to OmcZ as well, and yet the discrepancy remains at least ∼10^3^-fold after correcting for this observation. The discrepancies with experiment likely reflect additional physical artifacts of the solid-state measurements.

The simulations^30, 71, 73, 74^ were performed for a freely-floating protein in a bulk aqueous electrolyte with no externally applied bias (ET conditions), whereas the electrical measurements^3, 29, 30^ were conducted on bear Au electrode-adsorbed and air-dried protein exposed to an electric field unscreened by mobile ions (ETp conditions), and in the case of conducting probe atomic force microscopy, crushed under 10–50 nN of force.^29, 30^ To illustrate how different conditions change the nature of electrical conduction, biological ET in cytochromes requires redox activity, whereas ETp through cytochromes does not.^87, 110-112, 124^ It remains an open question as to what, if anything, about biological ET can be learned from solid-state ETp.^125^

Furthermore, forces of that 10-50 nN magnitude used in the experiments are known to mechanically deform much more structured proteins (*e.g.*, azurin, plastocyanin, and cytochrome *c*),^126-131^ and change the ETp mechanism; for example, by changing the packing density of the protein. More force can increase the packing density, which in turn promotes the transmission of electrons. The force needed to obtain a stable current from the ostensibly more conductive OmcZ filament^29^ was reported to be 5-fold larger than for OmcS^30^ (50 vs. 10 nN). In this context, the experimental procedure of measuring the length dependence of the resistance by having the current pass through previously crushed segments of the filament^29, 30^ may produce artifacts.

In general, the cytochrome filaments are probably more responsive to compressional force than previously studied well-structured proteins because ≥50% of the secondary structure of cytochrome filaments consists of flexible turns and loops.^3, 23-27^ OmcZ is probably more susceptible than OmcS because there is much less unstructured protein ‘insulation’ to provide a physical buffer between the AFM ‘tip’ and the heme chain.

The compressional crushing, as well as the effects of electrode adsorption and dehydration in the experiments have not yet been considered in computations because they are difficult to model,^112^ as well as being experimental artifacts with no bearing on the physiological redox process. The uncharacterized and uncontrolled level of dehydration in the experiments is important because Nguyen and co-workers^45^ have shown a ∼10^5^-fold variation in the conductivity of *G. sulfurreducens* biofilms with the level of hydration.

##### 3.1.5.2. No Structural Basis for near ‘Speed Limit’-breaking Electron Transfer Rates in OmcZ

The 2 × 10^12^ s^-1^ k_eff_for OmcZ is nearly at the non-adiabatic ‘speed limit’ of 10^13^ s^-1^ proposed by Gray and Winkler.^123^ The ‘speed limit’ was estimated by extrapolating the distance dependence of the rate for Cu(I) → Ru(III) electron transfer in Ru-labeled azurins to a metal-to-metal van der Waals contact distance of 3 Å. However, the metal-to-metal distance in all known cytochrome filaments is 9–11 Å and the heme-to-heme electronic couplings are weak (< k_b_T at T = 298 K). Furthermore, as discussed above, k_eff_is a *lower* bound estimate for the rate-limiting step in a heterogeneous chain with a maximal ρ, and neglects the siphoning-off of electrons onto the branching heme in OmcZ. Even if the effective rate is ∼10^2^-smaller because of the number of hemes coupled to the electrodes, the experimental rate is still 10^3^-fold faster than expected for a rate-limiting T-stacked heme pair in the structure.

Based on the conserved heme packing arrangements, as well as the largely universal energetic parameter ranges (Section 3.1.1) and kinetic rate constants (Section 3.1.2) for those heme packing topologies, there is no structural basis for 10^12^ s^1^ rates in a multi-heme protein with weak heme-to-heme couplings. The ∼10^13^ s^-1^ k_eff_ in GmP is even more unreasonable for a cytochrome.

### 3.2. Measured Conductivity Dependencies are Unphysical

In the following sub-sections, the reported dependence of filament conductivity on pili variants, crystalline heme packing, carbon nanotube-like charge propagation, temperature, and pH are discussed in Sections 3.2.1 – 3.2.5. Because a large structural change has been proposed to account for the pH-dependence, Section 3.2.6 discusses discrepancies between structural characterization methods.

#### 3.2.1. The Conductivity-Aromatic Density Correlation in Pili was Overstated and Now is Meaningless

If only cytochrome-based filaments exist, the prior sections demonstrate that the ∼10^7^-fold variation in conductivity reported by Malvankar and co-workers^18-20, 22^ is lacking in a structural basis, physiologically irrelevant, and for the largest values, physically implausible. Where does that leave the correlation Malvankar and co-workers reported between the measured conductivities and the aromatic density in the pilus?^22^

Before the CryoEM structures of cytochrome filaments, Malvankar and co-workers’ hypothesized that electrical conductivity should correlate with aromatic density if the MLP hypothesis was correct.^22^ This idea was tested by plotting (Figure 5, black) the logarithm of the measured conductivities for the putative GsAro5, GuP, GsP, GsW51W57, and GmP filaments as a function of the fractional composition of aromatic residues in the primary sequences of these filaments, dubbed aromatic residue density (ARD); note that GsAro5 was not included in the original figure of Ref. 22.

**Figure 5.**
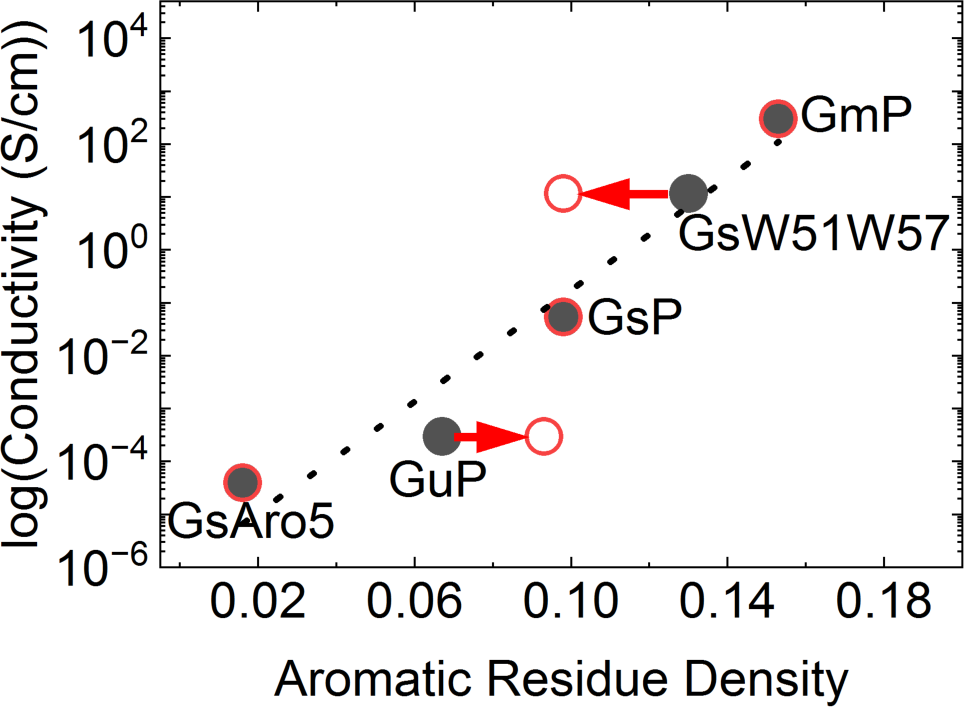
Correlation between the logarithm of the measured conductivity for filaments with the aromatic residue density of the supposed primary sequence of their building block. The five filaments were thought to be the *G. sulfurreducens* pilus with five aromatic residues (F24, Y27, Y32, F51, and Y57) mutated to Ala (GsAro5), the heterologously expressed *G. uraniireducens* pilus (GuP), the wild-type *G. sulfurreducens* pilus (GsP), the wild-type *G. sulfurreducens* pilus with Phe-51 and Tyr-57 mutated to Trp (GsW51W57), and the heterologously expressed *G. metallireducens* pilus (GmP). See the *Sequences of Putative Pili Used in the Conductivity-versus-Aromatic Density Correlation* section in the Supporting Information for more details.

The definition of ARD as the fractional aromatic composition was somewhat misleading because density is not a one-dimensional descriptor. The term “density” connotes packing of matter in some volume, but no structural information about the three-dimensional density of aromatic groups in the examined filaments was (or still is) available.

Note, too, that if the filaments are actually cytochromes, as now argued,^3, 29^ the correlation is between the conductivity in 5 different cytochromes and the aromatic residue density in the pilus. In a fortuitous series of pleotropic effects, somehow genetically introducing pili variants would have had to yield different cytochrome filaments whose conductivities varied in precisely the way expected by varying the aromatic density in the pilus. Another possibility is that the variation reflects uncontrolled variables in the experimental conditions.^45^ Given the foregoing discussion, the possibility that the ∼10^7^-fold variation is an experimental artifact may seem more plausible than supposing such a variation in conductivity can somehow be realized within a multi-heme architecture.

The conductivity-versus-aromatic density relationship shown in black on Figure 5 is linear (R^2^ = 0.9187), but there are several problems with the analysis, some of which pre-date the discovery of the CryoEM structures of GsP^28^ and cytochrome filaments.^3, 23-27^

##### 3.2.1.1. Problems with the Original Analysis

Under the assumption that the GsP filament was only composed of the PilA-N protein, 6 of 61 residues were aromatic (Phe, Tyr, His, or Trp), meaning that the ARD was 0.098. This filament had a conductivity of 4.6 × 10^-2^S/cm at pH 7.^18^ Substitution of Trp for two of those aromatic residues (Phe-51 and Tyr-57) gave GsW51W57, which had a larger ARD (0.13) and larger conductivity (1.0 × 10^1^ S/cm), in agreement with the MLP hypothesis. The ARD was higher for GsW51W57 versus GsP because the indole sidechain that replaced each phenol or tyrosyl ring is composed of two aromatic rings (benzene + pyrrole) and therefore was counted as two aromatic residues. However, an indole is not composed of independent benzene and pyrrole rings; the rings are fused into a single aromatic system. Counting each indole as two aromatic residues was not correct.^132^ The replacement of two aromatic residues (Phe-51 and Tyr-57) by two aromatic residues (W51 and W57) should not change the ARD, meaning that the 10^3^-fold variation between GsP and GsW51W57 was *never* genuinely explained by the MLP hypothesis.

GmP had a still larger ARD of 0.15 and conductivity of 2.8 × 10^2^S/cm, which, again, agreed with the MLP hypothesis. GuP had a lower ARD and lower conductivity, as anticipated by the MLP hypothesis. The ARD of GuP was reported to be 0.067, but the present author counts 18 of 193 residues as being aromatic (see the *Sequences of Putative Pili Used in the Conductivity-versus-Aromatic Density Correlation* section of the Supporting Information), meaning that the ARD should be 0.093. The conductivity of GuP was 3 × 10^-4^S/cm. Finally, GsAro5 had the lowest ARD (0.016) and the lowest conductivity (4 × 10^-5^ S/cm).

The original analysis showing a linear conductivity-versus-aromatic density correlation for putative pili seems to have been flawed because (1) the number of aromatic residues in GuP was under-counted and (2) each introduced Trp in GsW51W57 was double counted. When correcting for these issues (red arrows on Figure 5), the correlation coefficient falls from 0.9187 to 0.5977, thereby providing much weaker support for the MLP hypothesis.

##### 3.2.1.2. New Problems in Light of CryoEM Pili and Cytochrome Filament Structures

Malvankar and co-workers have since reported that GsP is a supramolecular assembly of heterodimers containing the PilA-N and PilA-C proteins, instead of the previously believed assembly of only PilA-N proteins (known at the time as PilA).^28^ It is not known how the F24A- 27A-Y32A-F51A-Y57A and F51W-F57W mutants of PilA-N change the assembly or morphology of the pilus. The compositions and structures of the heterologously expressed GuP and GmP are also not known. It is not known, or it is debated whether wild-type *G. sulfurreducens* expresses any of these filaments extracellularly.^28, 133^ If the filaments are cytochromes, with OmcS and OmcZ being GsP and GsW51W57, then the identity of the putative GsAro5, GuP, and GmP filaments is still unknown. Confusingly, the *Geobacter* strain that produces the putative GsAro5 filament was recently used by Malvankar and co-workers to obtain OmcS.^29^ Does this strain produce multiple filaments and if so, under what conditions can GsAro5 or OmcS be isolated? Or were GsP and GsAro5 the same filament measured under inconsistent experimental conditions?

Since the identity and composition of the filaments were and are still not known, the ARD is meaningless. The correlation rested on the *assumption* that the filament identity was a fact, not a questionable interpretation, or an assumption in its own right.

#### 3.2.2. Crystalline-like Heme Packing does not Contribute to Electron Delocalization

X-ray diffraction (XRD) analysis of pili^9, 17^/cytochrome^29^ preparations were reported by Malvankar and co-workers to show a sharp peak at 25°, corresponding to a *d*-spacing of ∼3.5 Å. The peak was reported to be more intense for GsW51W57/OmcZ than GsP/OmcS,^29^ and in both cases to increase in intensity upon acidification of the filaments.^17, 29^ Curiously, Malvankar and co-workers previously reported that the XRD peak disappeared for the putative GsAro5 pilus,^17^ but now report that the filament from the Gs*Aro5* strain of *Geobacter* produces the OmcS filament that shows a 3.5 Å spacing.^29^

The peak was reported to be “indicative of π-orbital overlap and charge delocalization.”^10^ Originally this π-delocalization was attributed to π-π stacking of aromatic groups in pili,^9, 17^ and now with the CryoEM structures of cytochrome filaments in hand, it is said to correspond to the face-to-face π-π stacking of heme cofactors. ^29^

This hypothesis can be tested in a straightforward way by simulating the XRD pattern expected from slipped- and/or T-stacked diheme model complexes^104^ in the Cambridge Crystallographic Data Center (CCDC) collection. The simulated XRD pattern for an ethane-bridged, bis-pyridine-3-carbonitrile ligated pair of (octaethylporphyrinato)iron(II) complexes (CCDC ID 1824039) has a slip-stacked arrangement of the macrocycles, and relatively weak intensity features in the 20-30° range that do not change if the ethane bridge and one of the porphyrins is deleted (Figure 6, *top*). The same diheme but with pyridine-4-carbonitrile axial ligands (CCDC ID 1824035) adopts T-stacked configurations of the diheme pairs in the crystal. The simulated XRD pattern for this crystal also shows relatively weak peaks that are invariant to whether one porphyrin (monomer) or a slip-stacked pair (dimer) is present (Figure 6, *bottom*). The T-stacking of two slip-stack pairs (tetramer) causes an increase in intensity.

**Figure 6.**
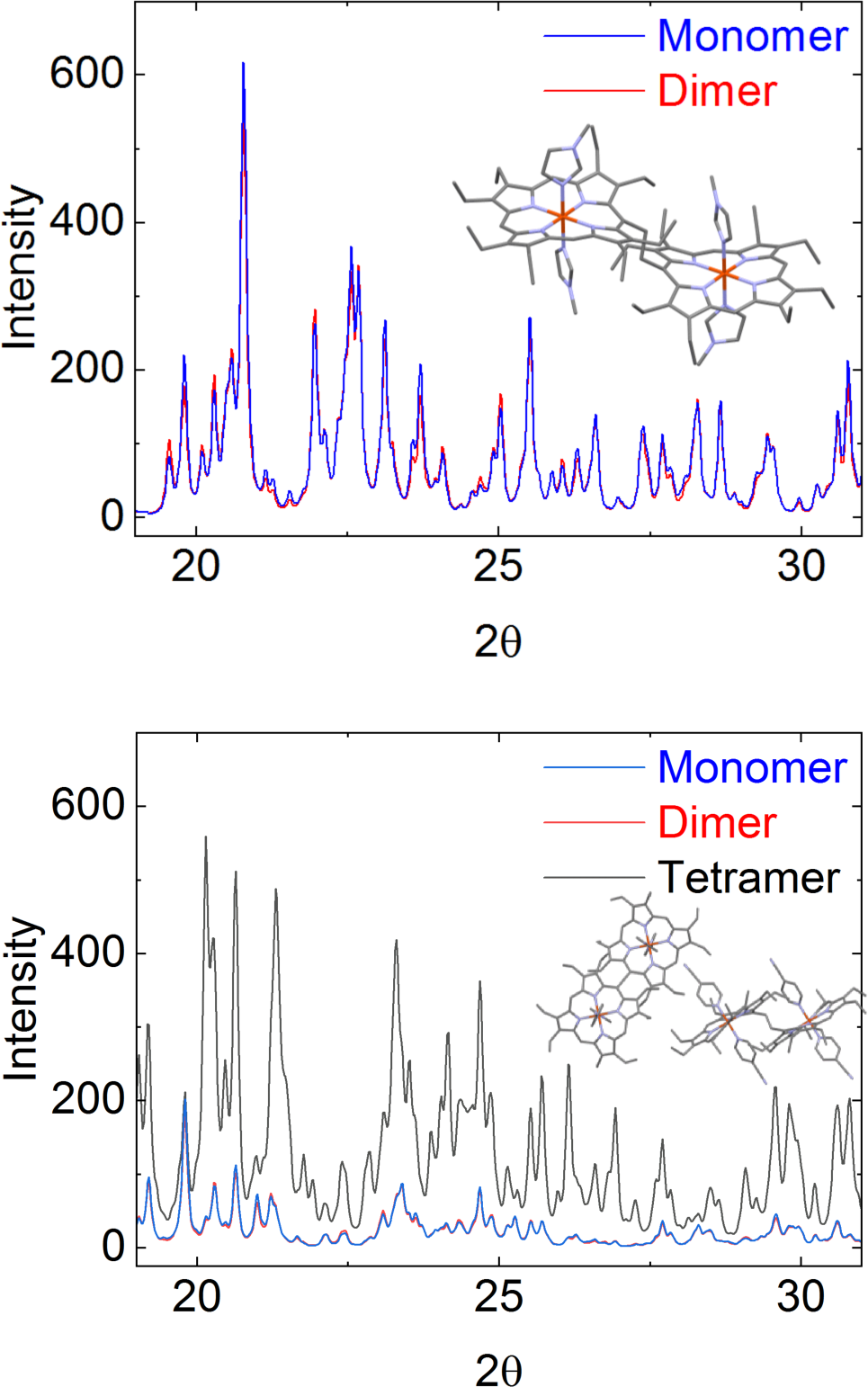
Simulated XRD powder patterns for model diheme systems in the CCDC collection.^104^ An ethane-bridged pair of (octaethylporphyrinato)iron(II) complexes with (*top*) pyridine-3- carbonitrile ligands (CCDC ID 1824039) and (*bottom*) pyridine-4-carbonitrile ligands (CCDC ID 1824039). The dihemes in the latter crystal pack in T-stacked configuration with respect to each other in the unit cell.

These results suggest that increased intensity can result from T-stacked, but not slip-stacked heme pairs. Since T-stacked heme pairs have the weakest electronic coupling of the two heme packing geometries, it is difficult to reconcile these results with Malvankar and co-workers’ interpretation that increased intensity of the peak from GsP/OmcS to GsW51W57/OmcZ or pH 10.5 to 2 for either filament indicates improved crystallinity through π-π stacking, which increases the effective conjugation length and yields a longer mean free path for electrons to enhance conductivity.^9, 17, 29^ This picture belongs to a synthetic organic metal, (or the formerly supposed metallic-like pilus), not a redox conductor with sub-thermal energy electronic couplings between charge relaying sites.^10, 74^ As discussed above, the electron transfer rates in neither Omc- S nor Z are large enough to satisfy a band theory transport description.

#### 3.2.3. Carbon Nanotube-like Charge Propagation may be an Experimental Artifact

Malvankar and co-workers “directly examined charge flow and distribution along individual native pili” and reported that holes (but not electrons) delocalized rapidly for microns along individual GsP filaments and accumulated in cytochrome-like globules on the pili. How is this observation consistent with the filament being polymerized OmcS and what, then, are the cytochrome-like globules (purification artifacts)? It was also observed that the mobile charge density was constant upon cooling and increased by 10^2^-fold upon acidification.^15^ All these observations were said to be carbon nanotube-like and suggested conductivity via delocalized charge instead of the redox conduction mechanism typically found in biology.^15^

Extensive charge delocalization is inconsistent with the CryoEM-resolved cytochrome filaments^3, 23-27^ and computations of their electronic structures,^74, 81^ that indicate mobile charges are localized in spatially well-defined heme-centered molecular orbitals under physiological (not necessarily solid-state) conditions. The experimental results may be an artifact of the ±10 V bias used in the electrostatic force microscopy experiments,^15^ which accessed electronic excited states completely forbidden to biology, or some other structure than the known cytochrome filaments may have been interrogated.

As noted in other sections, GsAro5 was reported to not show charge propagation, in contrast to GsP,^15^ but now the filaments harvested from the *GsAro5* strain is somehow OmcS, which showed extensive charge propagation. What filament was produced by the *GsAro5* strain that previously, but not now,^29^ showed properties distinctly different from OmcS and that nicely confirmed the MLP hypothesis? Did such a filament ever exist?

#### 3.2.4. An Erroneous Model Used to Explain Temperature Dependent Conductivity

Malvankar and co-workers reported that the conductivity of biofilms^9^ and films of pili^9^ (Figure 7, brown squares and circles, respectively) exponentially increased upon cooling down to a cross-over temperature close to the freezing point of water, below which the conductivity exponentially decreased upon further cooling. In the case of films of pili filaments, the temperature dependence of conductivity was 10^4^-fold smaller, the crossover temperature was lower, and the activation energy barrier was smaller— “all indicating reduced disorder and improved metallic nature.”^9^

**Figure 7.**
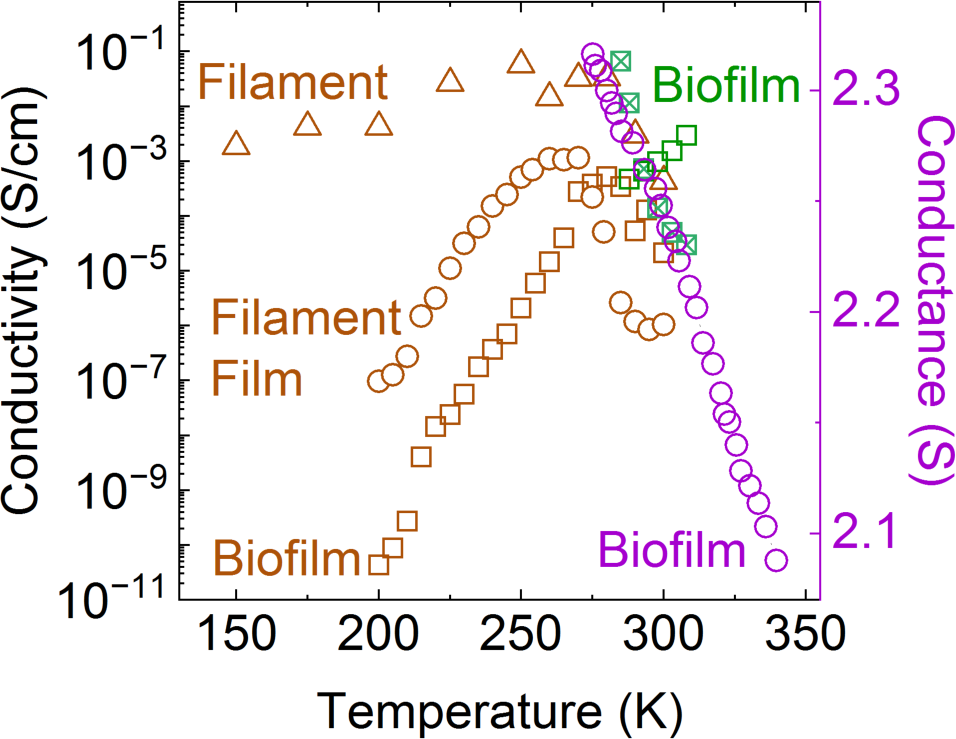
Previously reported temperature dependent conductivities for *G. sulfurreducens* biofilms (brown squares), films of filaments (brown circles), and individual filaments (brown triangles) according to Malvankar and co-workers.^9, 30^ Also shown are the temperature dependent conductance for films of filaments reported by Hochbaum and co-workers (purple circles),^58^ and the conductivity reported by Nguyen and co-workers under conditions of constant ambient water content or the relative humidity (green squares with and without inscribed X, respectively).^45^ Note the very different scale ranges on the left- and right-hand side vertical axes.

Malvankar argued that the bell-shaped temperature dependence was not consistent with cytochromes for which the expected redox conduction mechanism follows Arrhenius-type kinetics. The bell-shaped temperature dependence was instead consistent with delocalized electrons that move freely without thermal activation and experience reduced phonon scattering upon cooling, but encounter disorder, or localizing traps at temperatures below the crossover point, as in synthetic organic metals.^134^

Other laboratories at the time were not able to reproduce the cooling-induced exponential increase-then-decrease in conductivity over two orders of magnitude for either *G. sulfurreducens* biofilms or filaments. Hochbaum and co-workers^58^ found only a monotonic increase in conductance with a change in slope around 295 K and an overall change in conductivity down to 275 K of only 10% (Figure 7, purple; note the very different scale ranges on the left-versus right-hand sides of the figure). Reguera and co-workers found a decrease in differential transversal conductance upon cooling.^59^ Nguyen and co-workers^45^ showed that the conductivity of *G. sulfurreducens* biofilms increased or decreased upon cooling (Figure 7, green squares with and without an inscribed X), respectively, if the ambient water content or the relative humidity is kept constant. Both dependencies result from how the hydration state effects the mobility of ions coupled to redox reactions.

Malvankar and co-workers have subsequently reported^30^ that this same temperature dependence belongs to, instead of refutes the involvement of the OmcS cytochrome in conductivity (Figure 7, brown triangles). Specifically, Malvankar and co-workers (including the present author) interpreted the *anti*-Arrhenius (*i.e.*, increase in conductivity upon cooling) part of the temperature dependence in terms of a cooling-induced “massive restructuring”^30^ of hydrogen-bonds that changes the free energy landscape for electron transfer.

The level of humidity in the experiments reported by Malvankar and co-workers was never quantified, characterized, or controlled. Consistent with the Nguyen and co-workers’ observations, the report on OmcS^30^ presents “representative data” because the conductivity varied so much between replicates that averaging over the trials would not have produced a meaningful temperature trend.

Given the CryoEM structure of OmcS, a computational effort to validate the experimental temperature dependence in terms of redox conduction was reported by Malvankar and co-workers.^30^ The present author previously pointed out (above and elsewhere^73^) that the unphysically negative—and experimentally false (Figure 2)—potential at -0.521 V versus SHE at 310 K computed by Dahl *et al.*^30^ created a large (∼|0.44| eV) free energy difference that was larger than any free energy difference computed at 270 K. The result was a severe underestimation of redox conductivity at 310 K, and the *apparent* agreement with experiment regarding increased conductivity at 270 K, thereby validating the experimental results on fictitious grounds.

The dissertation with the spectroelectrochemical data^101^ that overturns the model pre-dated the Dahl *et al.* publication^30^ from the same (Malvankar) laboratory by two years, and the spectroelectrochemical measurements were reproduced by another member of the laboratory in that intervening time.^106^ In other words, a model predicated on knowingly false redox potentials, (Figure 2) but which validated the experimentally observed temperature dependent conductivity was published by Malvankar and co-workers.^30^ Meanwhile, a model that reproduced the spectroelectrochemical data (Figure 2) but did not support the observed temperature dependent conductivity was developed in the Malvankar laboratory by the present author and published by virtue of the independence afforded him by the National Institutes of Health Ruth L. Kirschstein Postdoctoral Fellowship.^73^ Hypothesis-refuting computations have not been published from the Malvankar laboratory.

It is also worth noting that several other theoretical errors plagued the Dahl *et al.* article, some of which are given below. These issues were raised by the present author who advised on the Dahl *et al.* work. These concerns were ignored, just like the concern expressed over the correctness of the redox potentials relative to spectroelectrochemistry. However, the present author did not appreciate how much any of these concerns changed the conclusions of the work until after the article was published.

1. A substantial change in H-bonding occupancy was misrepresented as a “massive restructuring” in H-bonding connectivity.^73^
2. An improper implementation of Kinetic Monte Carlo (KMC) was used to compute the charge diffusion coefficient by averaging displacements over the *number* of MC steps, which corresponded to unequally-sized time increments.^135, 136^ The KMC implementation was known *before* publication not to reproduce the results of a previously published implementation of the analytical Derrida formula,^82, 83^ which gave much lower charge diffusion coefficients.
3. An amalgam of forcefield parameters from different publications were used without validation for classical molecular dynamics: The CHARMM36 forcefield for proteins^137^ was paired with bonded parameters for the heme group developed for CHARMM27 by Autenrieth *et al.*^138^ and partial charges for the hemes and bonded cysteine and histidine residues from Barrozo *et al.*,^139^ which were not developed in accord with the prescriptions of the CHARMM forcefield. Because some heme parameters originated from an older version of the CHARMM forcefield than used for the protein, both generations of the forcefield were loaded into NAMD^140^ to perform the simulations, which retained unduplicated energy terms from both generations of the forcefield. The result was potentially an unvalidated admixture of different generations of the CHARMM forcefield to describe the protein.
4. Configurations from the ensemble with all hemes oxidized were used to compute electronic couplings between the charge donor and acceptor, but coupling does not occur near the minimum of the reactant potential energy surface. Configurations appropriate to the transition state region should be sampled, which can be approximated by assigning charges to the donor and acceptor hemes that are half-way between the reduced and oxidized states.^84,107^

#### 3.2.5. pH dependence explained with a Physically Unreasonable Mechanism

The conductivity of both GsP^9^ and GsW51W57^19^ were reported to increase upon acidification down to pH 2. Malvankar and co-workers argued that this result was inconsistent with conductivity via cytochromes because they denature at this low pH. The observation was instead more consistent with the ability of organic metals to be chemically doped (here by protons) to increase the charge carrier concentration, which enhances conductivity.^12^ This result was also said to support the idea that the charge carriers in pili were holes.

Malvankar and co-workers now attribute the same observations to the Omc- S and Z cytochromes,^29^ and argue that they become more, not less structured at pH 2. Infrared nanospectroscopy using scattering-type scanning near-field optical microscopy (IR s-SNOM) indicated^29^ a switch from ∼70% α-helical to ∼70% β-sheet content in both Omc- S and Z upon lowering the pH from 7 to 2. Circular dichroism (CD) data were interpreted as “consistent” with the s-NOM results, even though the reported analysis of the CD spectra indicated only 21 and 13% increases in β-sheet content, respectively. Assays with Thioflavin T (ThT) showed increased emission upon acidification, which *may* indicate greater β-sheet content. However, use of ThT under acidic conditions can also give false positives/negatives.^141^ The other employed spectroscopic techniques are also not unambiguous and the interpretations *presume* that the observations are entirely attributable to OmcZ in impure samples. More fundamentally, the geometrical constraints imposed on the polypeptide backbone by the two covalent thioether linkages and two coordinative His-Fe bonds to each heme make it seem doubtful that large changes in protein secondary structure can physically be accommodated in response to extreme (physiologically irrelevant for *G. sulfurreducens*) pH conditions.

Acidification was also attributed with causing a ∼1 nm reduction in filament diameter. The AFM-measured heights of Omc- S and Z decreased from 3.6 to 2.4 and 2.5 to 1.5 nm, respectively, from pH 7 to 2.^29^ Given that the diameter of a heme group is ∼1.5 nm,^72^ it is unclear how any protein can be present in the height measurement of OmcZ at pH 2, if OmcZ was in fact the filament being measured.

As an alternative to the structure-transition model, redox conduction can become more favorable with an increase in H^+^ concentration if the electron and proton transfers are coupled. The present author previously found that pKas in OmcS at physiological (5–7) pHs compensate for the change in net charge upon reduction by ∼50%.^73^ A likeness was also found between the large suppression in conductivity for OmcS upon deuteration and similarly large H/D kinetic isotope effects (KIEs) reported for other systems for which the observation was interpreted as proton-coupled electron transfer.^73^ The deuteration effect may also reflect the dynamical control of the environment over electron transfers,^142^ but it is unclear if the magnitude of this effect can account for the observed KIEs in OmcS.

It is worth noting that the attribution of the effect to pH is potentially misleading. The CD spectra reported for OmcS in a potassium phosphate pH 7 solution and a sodium citrate pH 2 solution are similar, but become much less so under solid-state measurement conditions.^29^ The so-called pH-dependence, which is already physiologically irrelevant at pH 2, may be an artifact of solid-state conditions.

#### 3.2.6. Spectroscopic Structural Characterizations inconsistent with CryoEM of OmcZ Filament

The spiraling heme chain at the core of all known cytochrome filaments is ‘insulated’ by a protein that has at least 50% of its 2° structure comprised of loops and turns.^3, 23-27^ The structures typically have 13-19% α-helical (the exception being F2KMU8 with 29%) and 3-6% β-sheet (the exceptions being F2KMU8 and A3MW92 with 15 and 18%, respectively) content.

Spectroscopic data published by Malvankar and co-workers^29^ strongly disagree with these secondary structural compositions from CryoEM analyses at pH 10.5. IR s-SNOM indicated that Omc- S and Z at pH 7 both have ∼70% α-helical content (instead of <20%), 10 or 29.6% β-sheet content (instead of ≤6%), respectively, and 20.8% or an undetectable percentage of coils/turns. CD spectra were interpreted as showing Omc- S and Z to have 65.55 vs. 38.7% α-helical and 15.75 versus 40.8% β-sheet content. Earlier CD structural analyses by others^143, 144^ were said to be consistent with these results, even though they indicated ∼10–13% α-helical content for either protein, and 23% β-sheet and 28% β-turn content for OmcZ. A simple explanation for these discrepancies is that the samples were insufficiently pure.

Malvankar and co-workers offered in the article^29^ a different explanation for the discrepancies between s-NOM (but not CD) and CryoEM analyses: Percentages from s-NOM are only quantitative in a comparative sense due to a particular sensitivity to C=O versus N-H stretching. If true, the finding of 19% more β-sheet content in Omc- Z versus S is still at odds with ∼6% β- sheet content in both CryoEM structures.^25, 26^ Moreover, Malvankar and co-workers state, at odds with their deposited structure (PDB 7LQ5), that the CryoEM model of OmcZ has 21 instead of 6% β-strands. It is also inappropriate to argue, as these authors do, that the total “β-strand and turn” content is larger in Omc- Z versus S, because turns do not qualify as regular secondary structure.

But the explanation for the s-NOM-versus-CryoEM discrepancy has other logical problems. First, it is strange because Malvankar and co-workers validated the s-NOM technique against the CryoEM structure of OmcS at pH 10.5. It appears that when the CryoEM structure is known beforehand, s-NOM agrees well with CryoEM; the CryoEM structures of OmcZ were only solved later.^25, 26^ Second, it is an inaccurate statement that demonstrates a misunderstanding about the technique. The original method article for s-NOM indicates that the technique “primarily probes molecular vibrations that oscillate perpendicular to the sample surface.”^145^ The sensitivity to C=O versus N-H stretching only pertains to the specific case studied in that article of membrane-embedded bacteriorhodopsin because of the orientation of the protein in the membrane. The orientation of OmcZ filaments on the surface, by contrast, was not controlled in s-NOM experiments. The take-away is that the secondary structure analysis from s-NOM is highly dependent on the orientation of the protein on the surface.

Another piece of characterization data from Malvankar and co-workers that is inconsistent with CryoEM analyses is the AFM-measured height for OmcZ. AFM indicated that the diameters of dried samples of Omc- S and Z originally in a pH 7 solution were respectively 3.6 and 2.5. The CryoEM structures at pH 10.5 indicate that the widest points on Om- S and Z are ∼4 and ∼5 nm, respectively.^24^ The difference likely reflects the effect of dehydration, but it should also be pointed out that the definition of the diameter is somewhat arbitrary from a structural point of view for a helical filament that is only approximately cylindrical. Whatever the reason for the discrepancy, the different diameters from AFM and CryoEM indicate difficulties with using AFM as a protein-identification technique before electrical characterizations.

## 4. Conclusions

The identity, structure, and *in vivo* mechanism of electrically conductive filaments expressed by *Geobacter sulfurreducens* have been intensely investigated and debated over the past 20 years. In that time, the hypothesis that a crystalloid array of stacked aromatic residues confers metallic-like conductivity to a pilus was advocated with utmost confidence. Experiments “definitively rules[d] out”^12^ and proved “physically impossible”^13^ the more traditional view of redox conduction through cytochromes, until, that is, CryoEM revealed the filaments to be cytochromes. Now, the filaments are claimed to have been cytochromes all along, and it is *assumed* that the characterization data that argued otherwise can simply be ascribed to them. Doubting the connection between structural and electrical characterizations, or the physiological relevance and physical plausibility of the latter, as done previously,^7, 44, 45, 73, 74^ may seem more sensible.

The present study posed the question: Are electrical characterizations consistent with the cytochrome structures of *Geobacter* ‘Nanowires?’ By trying to reconcile the characterization data in favor of a metallic-like pilus with the now-known CryoEM structures of cytochrome filaments, the following key conclusions were reached.

Electrons flow through cytochrome filaments (and multi-heme proteins more generally) in a succession of redox reactions for which the energetics are physically constrained (Section 3.1.1) and the kinetics within highly conserved heme packing geometries are largely independent of protein identity (Section 3.1.2). Computed heme-to-heme electron transfer rates for T- and slip-stacked heme pairs in cytochrome filaments agree, on average, within a factor of 10 of rates experimentally determined in other multi-heme proteins for the same heme packing geometries.

T-stacked heme pairs, which comprise nearly or exactly half of all heme pairs in cytochrome filaments are electronic coupling-constrained bottlenecks for electron transfer that set the rate-limiting reaction to the µs timescale, which is still much faster than typical ms enzymatic turnover. The rate-limiting step for respiration resides elsewhere in the cellular machinery. Tuning the conductivity of cytochromes over the reported ∼10^7^-fold range for filaments from *G. sulfurreducens* strains with pili variants seems both physically implausible and physiologically irrelevant if those filaments are supposed to be cytochromes.

Both the observation (Section 3.1.3) and magnitude (Sections 3.1.4–3.1.5) of conductivity for fully oxidized or fully reduced cytochrome filaments is inconsistent with physiologically relevant redox conduction through a chain of heme cofactors. The protein-limited flux for redox conduction through a 300-nm filament of both T- and slip-stacked heme pairs is predicted to be ∼0.1 pA (Section 3.1.4). This result implies that a *G. sulfurreducens* cell discharging ∼1 pA/s needs at least 10 filaments, which is similar to the experimental estimate that ≥20 filaments/cell are expressed.^1,146^

The experimental currents for the Omc- S and Z filaments at a physiologically relevant 0.1 V bias, however, are ∼10 pA and ∼10 nA, respectively. Some of the discrepancy is attributable to the experimental conditions of a dehydrated protein adsorbed on a bear Au-electrode that contacts ∼10^2^ hemes versus the simulation conditions of a fully hydrated and freely floating protein in electrolyte solution (Section 3.1.5.1). It is debatable what, if anything, about biological electron transfer can be learned from solid-state electron transport.^125^

The uncontrolled level of hydration in the experiments,^45^ as well as the use of compressional forces in conducting probe atomic force microscopy that are known to mechanically deform (*e.g.*, change the packing density) and alter the electron transport mechanism of more well-structured proteins^126-131^ also casts doubts on the meaningfulness of the electrical characterizations. A five-fold stronger force was needed to obtain a stable current from the ostensibly more conductive Omc- Z versus S (50 versus 10 nN) filament, and the length-dependent resistance experiments were performed such that the current must pass through previously crushed segments of the filament. Because Omc- S and Z differ in mechanical properties, even treating both proteins with the same force may produce different structural deformations and non-comparable conductivities. The reported conductivity for OmcZ *appears* physically implausible (Section 3.1.5.2), as it implies an effective electron transfer rate at least 10^3^-fold faster than expected for a rate-limiting T-stacked heme pair. If only one site on the filament is coupled to the electrode at each heterogeneous interface, the lower bound estimate for the effective rate exceeds the maximum rate for a hopping description and approaches the 10^13^ s^-1^ non-adiabatic ‘speed limit,’ which is non-sensical for weakly coupled metal centers separated by 9–11 Å. Given the CryoEM structure of OmcZ and the application of physiologically relevant redox conduction theory to it, as well as the knowledge that intra-filament electron transfer does not constrain cellular respiration, the reported conductivity for OmcZ seems to be an experimental artifact with no bearing on biology.

The correlation between a 10-million-fold variation in filament conductivity and percent protein aromatic composition, inappropriately called aromatic residue density, (Section 3.2.1) was flawed from the start by double-counting Trp residues and under-counting the aromatics in the *G. uraniireducens* pilus. The correlation is now rendered meaningless because the percent compositions do (or may) not correspond to the examined filaments, which are of uncertain identity. The proposition that the 5 filaments in the correlation are cytochromes with two of them being Omc- S and Z seems incredibly implausible: How can the introduction of 5 different genetic pili variants induce the pleotropic extracellular expression of 5 different cytochrome filaments that have conductivities that roughly correlate with the expectations of the metallic-like pilus hypothesis? And, given the energetic constraints and generic kinetic rate constants for electron transfers in highly conserved heme packing geometries, how can a ∼10^7^-fold variation be accommodated in polymerized multi-heme proteins?

Increased crystallinity (Section 3.2.2) detected by XRD with a *d*-spacing of ∼3.5 Å cannot rationalize the increased conductivity of Omc- Z versus S, or either filament at acidic versus neutral or basic pH because the description of delocalized electrons encountering less disorder is inappropriate for sub-thermal energy coupled redox centers. Carbon nanotube-like charge propagation (Section 3.2.3) over the entire filament for micron lengths is also inconsistent with the spatially well-defined and confined heme-centered molecular orbitals that mediate physiologically relevant redox conduction along the heme chain. The carbon nanotube-like behavior may be an artifact of the ±10 V biased used in the experiments, which access electronic states completely forbidden to biology.

The bell-shaped exponential increase-then-decrease in conductivity upon cooling (Section 3.2.4) for OmcS is inconsistent with the purely Arrhenius-style dependence expected for redox conduction (*i.e.*, thermally activated multi-step hopping). The temperature dependence could only be explained theoretically in the multi-step hopping framework with a model predicted on redox potentials known (but not publicly disclosed at the time) to be experimentally false, as well as a variety of other theoretical errors. A model that reproduces the spectroelectrochemical data for OmcS developed by the present author does not support the observed temperature dependence and attributed at least part of the discrepancy to changes in the microstructure of the protein hydration layers upon cooling.

The acidification-induced increase in conductivity (Section 3.2.5) may be attributable, in part, to coupled proton and electron transfers. However, spectroscopic and structural evidence also indicates that the so-called pH effect may be an artifact of the solid-state conditions under which the characterizations of the pH effect were conducted. Among those techniques, s-NOM detected a large pH-induced conversion to β-sheet structure but gave a secondary structure composition for OmcZ at higher pH (Section 3.2.6) that was wrong relative to CryoEM, both quantitatively and qualitatively in relation to OmcS. The other spectroscopies gave signals that are consistent with many other possibilities than the selected interpretation and presume that the observations are not attributable to impurities. More importantly, the proposed large-scale structural transition seems unphysical given the four (covalent or coordinative) bonds to each heme that constrain the polypeptide backbone conformation.

The sum total of the presented analysis is to answer the title question in the *negative*. A significant discrepancy currently exists—not between theory and experiment—but in terms of CryoEM structural versus functional characterizations of *Geobacter* ‘nanowires.’ Previously reported hallmarks of metallic-like conductivity are inappropriate and inconsistent with the CryoEM structures of cytochrome filaments. Meanwhile, the CryoEM structures, theoretical models, biological experiments, and kinetic analyses are all in agreement about the nature and rate of electron transfer in multi-heme architectures. The physiological relevance and/or physical plausibility of some experiments should be examined further.

## ASSOCIATED CONTENT

### Supporting Information

### The Supporting Information is available free of charge at

Heme packing metrical parameters, electron transfer energetics, assessment of active site polarizability on reorganization energy, comparison of rate constants, discussions of spectroelectrochemical data and intrinsic filament versus protein-electrode contact resistance, simulated currents, and protein sequences for conductivity-versus-aromatic density correlation.

## AUTHOR INFORMATION

### Corresponding Author

E-mail: * Phone: 475-225-6627

### Author Contributions

The research described herein was initiated and conducted solely by the corresponding author. Acknowledgements are given below to others who provided insightful advice on related questions.

### Notes

The author declares no competing financial interest.

## ACKNOWLEDGMENT

All calculations were performed using personal computing resources. The diffusion kinetic model was kindly provided by Fredrik Jansson. The multi-particle stead-state flux kinetic model was kindly provided by Blumberger and co-workers. Both models are integrated into the BioDC program developed by the present author.

