## Supporting Information for "Are Electrical Characterizations Consistent with the Cytochrome Structures of *Geobacter* ‘Nanowires’"

Table 1. Metrical Parameters for the heme centers in Omc- E, S, and Z from classical molecular dynamics simulations reported in Ref. 1.

| Heme Pair | Packing Designation | Edge-to-Edge Distance (Å) | Fe-to-Fe Distance (Å) | Heme Rotation (°) | Heme Plane Tilt (°) | His-His Rotation (°) |
|---|---|---|---|---|---|---|
| OmcE | | | | | | |
| 1↔2 | T | 6.1 ± 0.3 | 11.8 ± 0.4 | 136.3 ± 3.1 | 73.4 ± 4.5 | 32.1 ± 22.2, 16.2 ± 9.9 |
| 2↔3 | S | 3.7 ± 0.2 | 9.1 ± 0.3 | 178.0 ± 1.5 | 18.7 ± 4.4 | 16.2 ± 9.9, 20.0 ± 10.0 |
| 3↔4 | T | 6.0 ± 0.2 | 11.7 ± 0.4 | 145.5 ± 2.3 | 86.0 ± 3.2 | 20.0 ± 10.0, 72.7 ± 9.6 |
| 4↔1 | S | 4.4 ± 0.3 | 10.0 ± 3.3 | 172.0 ± 2.8 | 13.8 ± 4.9 | 72.7 ± 9.6, 23.1 ± 11.9 |
| OmcS | | | | | | |
| 1↔2 | T | 6.0 ± 0.2 | 12.6 ± 0.2 | 130.7 ± 3.0 | 69.1 ± 2.6 | 12.0 ± 9.1, 28.3 ± 11.2 |
| 2↔3 | S | 3.9 ± 0.2 | 9.4 ± 0.2 | 176.7 ± 1.7 | 11.0 ± 2.0 | 28.3 ± 11.2, 63.5 ± 10.1 |
| 3↔4 | T | 6.0 ± 0.2 | 11.3 ± 0.2 | 143.4 ± 2.2 | 75.2 ± 2.3 | 63.5 ± 10.1, 42.5 ± 11.2 |
| 4↔5 | S | 3.9 ± 0.2 | 9.5 ± 0.2 | 176.7 ± 1.8 | 8.0 ± 2.9 | 42.4 ± 11.2, 23,9 ± 12.0 |
| 5↔6 | T | 5.9 ± 0.2 | 11.3 ± 0.2 | 142.2 ± 2.3 | 82.2 ± 2.7 | 23.9 ± 12.0, 66.4 ± 10.9 |
| 6↔1' | S | 4.0 ± 0.2 | 9.7 ± 0.2 | 177.8 ± 1.6 | 7.7 ± 2.6 | 66.4 ± 10.9, 13.6 ± 8.2 |
| OmcZ | | | | | | |
| 1↔2 | T | 5.5 ± 0.1 | 11.4 ± 0.2 | 137.7 ± 3.3 | 68.8 ± 2.5 | 16.8 ± 8.9, 72.3 ± 10.2 |
| 2↔3 | S | 4.0 ± 0.2 | 9.8 ± 0.2 | 172.9 2.1 | 11.8 ± 2.4 | 72.3 ± 10.2, 45.9 ± 15.3 |
| 3↔8 | B | 5.3 ± 0.3 | 10.4 ± 0.3 | 96.5 ± 3.9 | 51.0 ± 3.5 | 45.9 ± 15.3, 40.9 ± 19.7 |
| 3↔4 | S | 4.2 ± 0.2 | 10.3 ± 0.2 | 56.5 ± 2.6 | 26.0 ± 3.1 | 45.9 ± 15.3, 12.5 ± 6.6 |
| 4↔5 | T | 4.9 ± 0.2 | 10.0 ± 0.3 | 172.4 ± 4.1 | 75.9 ± 4.2 | 12.5 ± 6.6, 16.4 ± 9.4 |
| 5↔6 | S | 4.0 ± 0.3 | 9.9 ± 0.4 | 160.8 ± 3.5 | 19.1 ± 3.9 | 16.4 ± 9.4, 57.2 ± 11.8 |
| 6↔7 | T | 5.3 ± 0.3 | 11.4 0.3 | 121.2 ± 4.0 | 75.4 ± 3.5 | 57.2 ± 11.8, 41.0 ± 17.9 |
| 7↔1' | S | 3.9 ± 0.2 | 9.4 ± 0.2 | 170.0 ± 2.4 | 23.3 ± 3.1 | 41.0 ± 17.9, 13,1 ± 7.2 |

Table S2. Electronic couplings computed for Omc- E, S, And Z surveyed from the literature.[1-3] The present author’s name (Guberman-Pfeffer) is abbreviated as G-P for conciseness.

| Author | Heme-to-Heme Electronic Coupling (meV) | | | | | | | | |
|---|---|---|---|---|---|---|---|---|---|
| | | | | | OmcE | | | | |
| | 4’↔1 | 1↔2 | 2↔3 | 3↔4 | 4↔1’’ | | | | |
| G-P. | 7.80 | 1.44 | 12.85 | 1.87 | 4.41 | | | | |
| | | | | | OmcS | | | | |
| | 6’↔1 | 1↔2 | 2↔3 | 3↔4 | 4↔5 | 5↔6 | 6↔1’ | | |
| G-P | 6.72 | 1.13 | 6.09 | 2.64 | 9.52 | 1.04 | 7.90 | | |
| Dahl et al. | | 8.93 | 15.9 | 3.48 | 12.4 | 5.68 | 8.76 | | |
| Jiang et al. | 3.94 | 1.54 | 9.35 | 2.12 | 6.07 | 1.18 | | | |
| | | | | | OmcZ | | | | |
| | 7’↔1 | 1↔2 | 2↔3 | 3↔4 | 4↔5 | 5↔6 | 6↔7 | 7↔1” | 3↔8 |
| G-P. | 9.83 | 2.04 | 3.19 | 7.15 | 4.06 | 5.21 | 2.32 | 4.71 | 2.43 |

Table S3. Reorganization energies (eV) computed for Omc- E, S, and Z surveyed from the literature.[1-3] The present author's name (Guberman-Pfeffer) is abbreviated as G-P for conciseness. G-P and Dahl *et al.* computed the reorganization energies using vertical energy gaps computed along classical molecular dynamics trajectories. Jiang *et al.* used an approximation to the Marcus continuum expression that parameterizes the static dielectric constant in terms of a linear dependence on the solvent accessible surface area, where the parameters were developed so that the reorganization energies reproduce those obtained from polarizable classical molecular dynamics.

| Author | Heme-to-Heme Electronic Coupling (meV) | | | | | | | | |
|---|---|---|---|---|---|---|---|---|---|
| OmcE | | | | | | | | | |
| | 4'↔1 | 1↔2 | 2↔3 | 3↔4 | 4↔1'' | | | | |
| G-P. | 0.688 | 0.879 | 0.481 | 0.780 | 0.778 | | | | |
| OmcS | | | | | | | | | |
| | 6'↔1 | 1↔2 | 2↔3 | 3↔4 | 4↔5 | 5↔6 | 6↔1' | | |
| G-P | 0.668 | 0.983 | 0.429 | 0.626 | 0.765 | 0.630 | 0.724 | | |
| Dahl et al. | | 0.605 | 0.578 | 0.658 | 0.829 | 0.819 | 0.675 | | |
| Jiang et al. | 0.66 | 0.86 | 0.64 | 0.71 | 0.60 | 0.78 | | | |
| OmcZ | | | | | | | | | |
| | 7'↔1 | 1↔2 | 2↔3 | 3↔4 | 4↔5 | 5↔6 | 6↔7 | 7↔1" | 3↔8 |
| G-P. | 0.593 | 0.556 | 0.785 | 0.892 | 0.781 | 0.758 | 0.680 | 0.624 | |

Table S4. Vertical energy gaps and reorganization energies (in eV) for the hemes of OmcE with self-consistently optimized electron densities in vacuum and the protein environment, or with the density frozen at the vacuum-optimized distribution while the heme is in the protein context. The data is reproduced from Ref. 1.

| Heme | Relaxed Density in Vacuum | | | Frozen Vacuum density In Protein Environment | | | Relaxed Density in Protein environment | | |
|---|---|---|---|---|---|---|---|---|---|
| # | -VEA | VIP | λ | -VEA | VIP | λ | -VEA | VIP | λ |
| 4’ | 4.881 ± 0.008 | 4.928 ± 0.009 | 0.023 | 3.250 ± 0.019 | 4.878 ± 0.018 | 0.814 | 3.239 ± 0.018 | 4.889 ± 0.019 | 0.825 |
| 1 | 4.874 ± 0.008 | 4.852 ± 0.008 | -0.011 | 2.961 ± 0.018 | 4.742 ± 0.020 | 0.891 | 3.013 ± 0.019 | 4.739 ± 0.020 | 0.863 |
| 2 | 4.869 ± 0.007 | 4.917± 0.009 | 0.024 | 3.008 ± 0.021 | 4.763 ± 0.019 | 0.878 | 3.064 ± 0.023 | 4.763 ± 0.020 | 0.850 |
| 3 | 4.808 ± 0.006 | 4.816 ± 0.009 | 0.004 | 3.084 ± 0.019 | 4.705 ± 0.021 | 0.811 | 3.135 ± 0.021 | 4.710 ± 0.020 | 0.788 |
| 4 | 4.860 ± 0.007 | 4.944 ± 0.009 | 0.042 | 3.168 ± 0.021 | 4.836 ± 0.040 | 0.834 | 3.181 ± 0.021 | 4.862 ± 0.018 | 0.841 |
| 1’’ | 4.882 ± 0.008 | 4.863 ±0.008 | -0.009 | 2.970 ± 0.024 | 4.871 ± 0.023 | 0.951 | 3.044 ± 0.024 | 4.865 ± 0.021 | 0.911 |

Table S5. Vertical energy gaps and reorganization energies (in eV) for the hemes of OmcS with self-consistently optimized electron densities in vacuum and the protein environment, or with the density frozen at the vacuum-optimized distribution while the heme is in the protein context. The data is reproduced from Ref. 1.

| | Relaxed Density in Vacuum | | | Frozen Vacuum density In Protein Environment | | | Relaxed Density in Protein environment | | |
|---|---|---|---|---|---|---|---|---|---|
| Heme # | -VEA | VIP | λ | -VEA | VIP | λ | -VEA | VIP | λ |
| 6’ | 4.949 ± 0.008 | 4.930 ± 0.007 | -0.010 | 3.449 ± 0.019 | 5.002 ± 0.017 | 0.777 | 3.464 ± 0.020 | 5.003 ± 0.016 | 0.770 |
| 1 | 4.888 ± 0.007 | 4.904 ± 0.008 | 0.008 | 3.326 ± 0.021 | 5.027 ± 0.019 | 0.851 | 3.372 ± 0.021 | 5.033 ± 0.020 | 0.831 |
| 2 | 4.900 ± 0.007 | 4.912 ± 0.008 | 0.006 | 3.327 ± 0.016 | 5.062 ± 0.018 | 0.868 | 3.329 ± 0.016 | 5.047 ± 0.017 | 0.859 |
| 3 | 4.834 ± 0.006 | 4.856 ± 0.006 | 0.011 | 3.530 ± 0.015 | 4.825 ± 0.016 | 0.648 | 3.555 ± 0.016 | 4.856 ± 0.015 | 0.651 |
| 4 | 4.943 ± 0.008 | 4.932 ± 0.008 | -0.005 | 3.344 ± 0.018 | 4.692 ± 0.018 | 0.674 | 3.398 ± 0.019 | 4.691 ± 0.018 | 0.647 |
| 5 | 4.870 ± 0.007 | 4.829 ± 0.007 | -0.021 | 3.342 ± 0.019 | 4.889 ± 0.018 | 0.774 | 3.333 ± 0.018 | 4.881 ± 0.018 | 0.774 |
| 6 | 4.932 ± 0.008 | 4.935 ± 0.009 | 0.001 | 3.392 ± 0.019 | 4.922 ± 0.018 | 0.765 | 3.421 ± 0.019 | 4.938 ± 0.019 | 0.759 |
| 1’’ | 4.869 ± 0.007 | 4.888 ± 0.007 | 0.010 | 3.334 ± 0.019 | 5.027 ± 0.018 | 0.847 | 3.356 ± 0.020 | 5.032 ± 0.018 | 0.838 |

Table S6. Vertical energy gaps and reorganization energies (in eV) for the hemes of OmcZ with self-consistently optimized electron densities in vacuum and the protein environment, or with the density frozen at the vacuum-optimized distribution while the heme is in the protein context. The data is reproduced from Ref. 1.

| | Relaxed Density in Vacuum | | | Frozen Vacuum density In Protein Environment | | | Relaxed Density in Protein environment | | |
|---|---|---|---|---|---|---|---|---|---|
| Heme # | -VEA | VIP | λ | -VEA | VIP | λ | -VEA | VIP | λ |
| 7' | 4.869 ± 0.007 | 4.879 ± 0.007 | 0.005 | 3.148 ± 0.021 | 5.126 ± 0.018 | 0.989 | 3.207 ± 0.022 | 5.146 ± 0.018 | 0.970 |
| 1 | 4.884 ± 0.008 | 4.877 ± 0.006 | -0.004 | 3.147 ± 0.022 | 4.797 ± 0.016 | 0.825 | 3.170 ± 0.024 | 4.806 ± 0.016 | 0.818 |
| 2 | 4.848 ± 0.007 | 4.862 ± 0.008 | 0.007 | 2.947 ± 0.019 | 4.815 ± 0.018 | 0.934 | 2.953 ± 0.020 | 4.821 ± 0.019 | 0.934 |
| 3 | 4.904 ± 0.007 | 4.912 ± 0.007 | 0.004 | 2.811 ± 0.020 | 4.924 ± 0.019 | 1.057 | 2.822 ± 0.021 | 4.913 ± 0.018 | 1.046 |
| 4 | 4.884 ± 0.007 | 4.856 ± 0.007 | -0.014 | 2.637 ± 0.019 | 4.721 ± 0.020 | 1.042 | 2.682 ± 0.020 | 4.728 ± 0.020 | 1.023 |
| 5 | 4.895 ± 0.008 | 4.883 ± 0.008 | -0.006 | 2.783 ± 0.018 | 4.696 ± 0.019 | 0.957 | 2.807 ± 0.018 | 4.692 ± 0.018 | 0.943 |
| 6 | 4.894 ± 0.008 | 4.892 ± 0.008 | -0.001 | 2.751 ± 0.020 | 4.824 ± 0.020 | 1.037 | 2.770 ± 0.020 | 4.820 ± 0.019 | 1.025 |
| 7 | 4.882 ± 0.008 | 4.885 ± 0.008 | 0.002 | 3.067 ± 0.018 | 5.008 ± 0.019 | 0.971 | 3.088 ± 0.018 | 5.019 ± 0.018 | 0.966 |
| 1" | 4.866 ± 0.006 | 4.870 ± 0.008 | 0.002 | 3.088 ± 0.017 | 4.810 ± 0.019 | 0.861 | 3.108 ± 0.018 | 4.813 ± 0.018 | 0.853 |
| 8 | 4.869 ± 0.007 | 4.853 ± 0.008 | -0.008 | 2.563 ± 0.021 | 4.883 ± 0.022 | 1.160 | 2.600 ± 0.020 | 4.879 ± 0.021 | 1.140 |

Table S7. Reaction free energies (eV) computed for Omc- E, S, and Z surveyed from the literature.[1-3] The present author's name (Guberman-Pfeffer) is abbreviated G-P for conciseness. G-P and Dahl *et al.* computed the reaction free energies using vertical energy gaps from quantum mechanical/molecular mechanical calculations at classical molecular dynamics-generated configuration (QM/MM2MD). However, the level of QM model chemistry, the number of evaluated configurations, and other technical aspects had important differences between the two studies. Jiang *et al.* approximated the reaction free energy as the redox-linked difference in electrostatic energy from Poisson-Boltzmann calculations.

| Author | Heme-to-Heme Electronic Coupling (meV) | | | | | | | | |
|---|---|---|---|---|---|---|---|---|---|
| | | | | | OmcE | | | | |
| | 4'↔1 | 1↔2 | 2↔3 | 3↔4 | 4↔1'' | | | | |
| G-P. | 0.188 | -0.038 | -0.009 | -0.099 | 0.067 | | | | |
| | | | | | OmcS | | | | |
| | 6'↔1 | 1↔2 | 2↔3 | 3↔4 | 4↔5 | 5↔6 | 6↔1' | | |
| G-P | 0.031 | 0.014 | -0.017 | 0.161 | -0.063 | -0.072 | -0.015 | | |
| Dahl et al. | | 0.018 | 0.209 | 0.186 | -0.440 | 0.111 | | | |
| Jiang et al. | 0.120 | -0.090 | 0.000 | 0.090 | -0.040 | -0.090 | | | |
| | | | | | OmcZ | | | | |
| | 7'↔1 | 1↔2 | 2↔3 | 3↔4 | 4↔5 | 5↔6 | 6↔7 | 7↔1'' | 3↔8 |
| G-P. | 0.189 | 0.101 | 0.019 | 0.163 | -0.044 | -0.046 | -0.258 | 0.092 | |

**Note on Interpreting Spectroelectrochemical Titration Curves**

The experimental curve in Figure 2 (*top*) of the main text reports on the macroscopic redox state $\left(S_x^y\right)$ of the OmcS filament as a function of solution potential. The subscripted x takes values from 0 to N, where N is the number of added reducing equivalence. Because the OmcS filament is a homopolymer of a hexa-heme protein and each heme undergoes a one-electron reduction, it may be assumed that there are $6 + 1$ or 7 macroscopic states in which 0, 1, 2, … 6 hemes are reduced. Implicit in this assumption is the idea that the redox properties of a subunit are independent of its position in the helical pitch of the filament (~4 subunits per pitch).

Each macroscopic state indexed by x is an ensemble of y microstates in which different combinations of hemes are reduced/oxidized.[4] The values of the superscripted y in the $S_x^y$ designation for the macroscopic redox state follows Pascal's triangle for the $x^{th}$ row: There is only one way to have all oxidized $(S_0^1)$ or all reduced $(S_6^1)$ hemes in OmcS, but there are 20 different combinations when 3-out-of-6 hemes are reduced $(S_6^{20})$.

The macroscopic redox potential for the $S_x \rightarrow S_{x+1}$ transition $(E_{x \rightarrow x+1})$ is related to the solution potential (E) by the Nernst equation (Eq. S1)

$$E = E_{x+1} - \frac{RT}{nF} \ln\left(\frac{[S_{x+1}]}{[S_x]}\right) \tag{S1}$$

where R, T, n, F, and $\left[S_{x/x+1}\right]$ are the gas constant, absolute temperature, number of transferred electrons, Faraday's constant, and the population of the $S_{x/x+1}$ state.

Under the approximation that a single microstate $(y = 1)$ dominates each macroscopic state $(x = 0, 1, 2, \ldots 6)$, $E_{x \rightarrow x+1}$ and $\left[S_{x/x+1}\right]$ in Eq. S1 can be replaced with the redox potential of an individual heme $(e_{x \rightarrow x+1})$ and the oxidized/reduced population of that heme $\left(\left[C_{ox/red}\right]\right)$. This approximation is only valid if redox cooperativities due to heme-heme interactions are negligible.

A Debye-Hückel shielded electrostatics model fitted to heme-heme interaction data for 17 multiheme proteins from five microbial genera[5, 6] predicts ~0.06 V shifts in $e_{x \rightarrow x+1}$ for hemes separated by Fe-to-Fe distances of ~11 Å (on average) as in OmcS.[6] An ethane-bridged complex of bis-4-methylimidzole (*i.e.*, histidine sidechain) ligated (octaethylporphyrinato)iron(III) molecules showed a -0.06 V shift in redox potential relative to the monomeric molecules.[7] The present author previously found that $e_{x \rightarrow x+1}$s differed by ≤0.07 V when the 6 hemes in the central subunit of an OmcS filament are titrated individually or simultaneously in Constant Redox and pH Molecular Dynamics (C(E,pH)MD)) simulations. Because computed $e_{x \rightarrow x+1}$ reasonably match spectroelectrochemically observed $E_{x \rightarrow x+1}$ (Table S8), the approximation seems warranted within the limited resolution of the experimental data. Also, no significant redox-linked conformational change was observed in the MD simulations, suggesting that the cause for order-of-magnitude stronger heme-heme interactions reported for the MtrF deca-heme protein was not operative in OmcS.[8]

Table S8. Comparison of spectroelectrochemically-derived and computed redox potentials. All values are in V vs. SHE. The parenthetical numbers indicate the heme to which the computed potential belong. The spectroelectrochemical data is reproduced from Ref. 9. The computed potentials from the present author (Guberman-Pfeffer) are reproduced from Ref. 10; those from Dahl *et al.* are reproduced from Ref. 3.

| | | Cathodic Direction | | Anodic Direction |
|---|---|---|---|---|
| | | Guberman-Pfeffer | Dahl *et al.* | Dahl et al. |
| Site | Experiment | QM/MM@MD | QM/MM@MD | QM/MM@MD |
| 1 | -0.070 | -0.108 ± 0.023 (#3) | -0.080 ± 0.06 (#6) | -0.421 ± 0.05 (#2) |
| 2 | -0.121 | -0.119 ± 0.027 (#1) | -0.108 ± 0.04 (#2) | -0.431 ± 0.06 (#6) |
| 3 | -0.138 | -0.128 ± 0.028 (#6) | -0.126 ± 0.05 (#3) | -0.491 ± 0.05 (#1) |
| 4 | -0.168 | -0.130 ± 0.023 (#2) | -0.191 ± 0.05 (#1) | -0.647 ± 0.05 (#4) |
| 5 | -0.220 | -0.214 ± 0.025 (#5) | -0.334 ± 0.04 (#4) | -0.727 ± 0.05 (#3) |
| 6 | -0.281 | -0.271 ± 0.027 (#4) | -0.521 ± 0.05 (#5) | -0.826 ± 0.05 (#5) |
| $E^{\circ}_{app}$ | -0.155 | -0.146 | -0.166 | -0.570 |

[a]The present study is chiefly concerned with the cathodic direction because most conductivity experiments are performed in the air-oxidized state. The computed anodic currents from Dahl *et al.* are also shown, however, because they too conflict with the spectroelectrochemical data that found a minimal (tens of mV at most) hysteresis.

Table S9. Heme-to-heme electron transfer rates for Omc- E, S, and Z surveyed from the literature.[1-3]

| | | Guberman-Pfeffer | | Jiang *et al.* | | Dahl *et al.* | |
|---|---|---|---|---|---|---|---|
| Heme Pair | Packing Designation | $k_{\rightarrow}$ | $k_{\leftarrow}$ | $k_{\rightarrow}$ | $k_{\leftarrow}$ | $k_{\rightarrow}$ | $k_{\leftarrow}$ |
| OmcE | | | | | | | |
| 4'↔1 | S | 2.5E+07 | 3.7E+10 | | | | |
| 1↔2 | T | 1.6E+07 | 3.7E+06 | | | | |
| 2↔3 | S | 4.6E+10 | 3.3E+10 | | | | |
| 3↔4 | T | 1.9E+08 | 4.1E+06 | | | | |
| 4↔1’’ | S | 5.2E+07 | 6.9E+08 | | | | |
| OmcS | | | | | | | |
| 6'↔1 | S | | | 5.0E+09 | 4.0E+07 | | |
| 1↔2 | T | 1.2E+06 | 2.1E+06 | 2.0E+06 | 5.0E+07 | 2.7E+10 | 1.1E+09 |
| 2↔3 | S | 2.1E+10 | 1.1E+10 | 4.0E+09 | 3.0E+09 | 9.0E+09 | 1.8E+10 |
| 3↔4 | T | 1.0E+07 | 5.2E+09 | 4.0E+08 | 2.0E+07 | 1.3E+07 | 4.2E+10 |
| 4↔5 | S | 3.4E+09 | 3.0E+08 | 1.0E+09 | 5.0E+09 | 2.7E+07 | 3.6E+10 |
| 5↔6 | T | 1.9E+08 | 1.2E+07 | 2.0E+06 | 7.0E+07 | 7.2E+10 | 2.9E+03 |
| 6↔1" | S | 1.5E+09 | 8.3E+08 | | | 9.2E+07 | 6.7E+09 |
| OmcZ | | | | | | | |
| 7'↔1 | S | 9.8E+07 | 1.5E+11 | | | | |
| 1↔2 | T | 5.1E+07 | 2.5E+09 | | | | |
| 2↔3 | S | 6.7E+07 | 1.4E+08 | | | | |
| 3↔4 | S | 5.2E+06 | 2.9E+09 | | | | |
| 4↔5 | T | 3.7E+08 | 6.8E+07 | | | | |
| 5↔6 | S | 8.1E+08 | 1.4E+08 | | | | |
| 6↔7 | T | 8.7E+09 | 4.0E+05 | | | | |
| 7↔1’’ | S | 1.7E+08 | 5.9E+09 | | | | |

Table S10. Comparison of measured and computed heme-to-heme electron transfer rates in various multi-heme cytochromes[1-3, 11, 12]

| | T-stacked | | | Slip-stacked | | |
|---|---|---|---|---|---|---|
| | Min. | Max. | Avg. | Min. | Max. | Avg. |
| Experiment | | | | | | |
| van Wonderen | 8.7E+6 | 1.3E+8 | 8.4E+7 | 3.3E+6 | 1.1E+10 | 2.8E+9 |
| OmcE | | | | | | |
| Guberman-Pfeffer | 3.7E+06 | 1.9E+08 | 5.3E+07 | 2.5E+07 | 4.6E+10 | 1.95E+10 |
| OmcS | | | | | | |
| Guberman-Pfeffer | 1.2E+06 | 5.2E+09 | 9.0E+08 | 3.0E+08 | 2.1E+10 | 6.34E+09 |
| Jiang et al. | 2.0E+06 | 4.0E+08 | 9.1E+07 | 4.0E+07 | 5.0E+09 | 3.01E+09 |
| Dahl et al. | 2.9E+03 | 7.2E+10 | 2.4E+10 | 2.7E+07 | 3.6E+10 | 1.16E+10 |
| OmcZ | | | | | | |
| Guberman-Pfeffer | 4.00E+05 | 8.70E+09 | 1.95E+09 | 5.20E+06 | 1.50E+11 | 1.60E+10 |

Table 11. Simulated steady-state current through a chain of all T-stacked, all slip-stacked, alternating T- and slip-stacked (as in OmcS, OmcE, A3MW92, and F2KMU8), or mixed T- and slip-stacked (as in OmcZ) heme groups as a function of the number of heme-to-heme electron transfer steps. The forward and backward rates constants were assumed to be $1 \times 10^8$ and $1 \times 10^9$ $s^{-1}$ for T- and slip-stacked pairs, respectively.

| Step # | All T-stacked | All Slip-stacked | Alternating | Mixed |
|---|---|---|---|---|
| 1 | 16.02 | 159.88 | 16.02 | 16.02 |
| 2 | 8.01 | 80.10 | 14.56 | 14.56 |
| 3 | 5.33 | 53.35 | 7.63 | 13.34 |
| 4 | 4.01 | 40.05 | 7.29 | 7.29 |
| 5 | 3.20 | 32.04 | 5.00 | 6.97 |
| 6 | 2.68 | 26.75 | 4.85 | 4.85 |
| 7 | 2.29 | 22.91 | 3.73 | 4.71 |
| 8 | 2.00 | 20.03 | 3.64 | 3.64 |
| 9 | 1.78 | 17.78 | 2.96 | 3.56 |
| 10 | 1.60 | 16.02 | 2.92 | 3.48 |
| 11 | 1.46 | 14.56 | 2.47 | 2.87 |
| 12 | 1.33 | 13.34 | 2.44 | 2.80 |
| 13 | 1.23 | 12.32 | 2.11 | 2.39 |
| 14 | 1.14 | 11.44 | 2.09 | 2.35 |
| 15 | 1.07 | 10.69 | 1.84 | 2.05 |
| 16 | 1.00 | 10.01 | 1.83 | 2.03 |
| 17 | 0.94 | 9.42 | 1.63 | 2.00 |

### Note on Intrinsic versus Contact Resistance

The contact resistance ($R_{cnt}$) was reported to be $\sim 1 \times 10^9$ and $\sim 1 \times 10^6$ for Omc- S[3] and Z[13] respectively. The length of either filament needed so that $R_{cnt} \ll R_{int}$ can be determined by multiplying the known length of the filament that bridged the electrodes $(3.0 \times 10^{-5}$ cm$)$ by the resistance-per-subunit $\left(\frac{R_{int}}{n_{sub}}\right)$, where $n_{sub}$ is the number of subunits.

Using standard relationships for conductivity, conductance, and resistances, and the fact that the length of the filament is an integer multiple of the length of a subunit, the resistance-per-subunit is given by Eq. S1.

$$\frac{R_{int}}{n_{sub}} = \frac{L_{sub}}{\sigma_{int} A} \tag{S1}$$

In this expression, $L_{sub}$ is the length of the subunit in the CryoEM model $(4.7 \times 10^{-7}$ or $5.8 \times 10^{-7}$ cm$)$; $\sigma_{int}$ is the intrinsic conductivity of the filament $(2.4 \times 10^{-2}$ or $3.0 \times 10^{-1}$ S$)$, and A is the cross-sectional area of the conductor, which is approximated as $\pi r_c^2$ for a cylinder, where $r_c$ is half the filament height measured by AFM $(2.0 \times 10^{-7}$ or $1.3 \times 10^{-7}$ cm$)$. The numerical values are given as (OmcS or OmcZ).

$\frac{R_{int}}{n_{sub}}$ comes out to $1.6 \times 10^8$ (OmcS) and $3.9 \times 10^5$ (OmcZ) $\Omega$, implying that the filaments need to be much longer than ~6 subunits (~30 nm) for OmcS and ~3 subunits (~15 nm) for OmcZ for the filament resistance to exceed the contact resistance and for the current to be protein-limited.

**Sequences of Putative Pili used in the Conductivity-versus-Aromatic Density Correlation***

```
  GsAro5: FTLIELLIVVAIIGILAAIAIPQASAARVKAANSAASSDLRNLKTALESAAADDQTAPPES------
     GuP: FTLIELLIVVAIIGILAAIAIPQFSKYRIQGFNASGNSDLKNIRTSQESLYAEWQHYGLTQGL----
     GsP: FTLIELLIVVAIIGILAAIAIPQFSAYRVKAYNSAASSDLRNLKTALESAFADDQTYPPES------
GsW51W57: FTLIELLIVVAIIGILAAIAIPQFSAYRVKAYNSAASSDLRNLKTALESAWADDQTWPPES------
     GmP: FTLIELLIVVAIIGILAAIAIPQFAAYRQKAFNSAAESDLKNTKTNLESYYSEHQFYPN--------


  GsAro5: -------------------------------------------------------------------
     GuP: ATVAGLPGAGKWGVGALVTPTAALPVCIITTDDNNLVPRGLQIPVGNNVTAMATTAAAGAGDGGSYT
     GsP: -------------------------------------------------------------------
GsW51W57: -------------------------------------------------------------------
     GmP: -------------------------------------------------------------------

  GsAro5: -------------------------------------------------------------------
     GuP: LAAKHLQGDVIFAADSDSTANYKMTFAAPATGLNAGYPLTAAFVPVSVNNVDDYQALPNWVKM----
     GsP: -------------------------------------------------------------------
GsW51W57: -------------------------------------------------------------------
     GmP: -------------------------------------------------------------------
```

*Note that GsP is now known to be a heterodimer of the PilA-N sequence (shown here) and PilA-C (not shown), but the filament examined in either case is now argued to be the OmcS cytochrome filament. The same disclaimer applies to GsW51W57, which is now argued to be the OmcZ cytochrome filament. Whether GsAro5, GuP, or GmP are only composed of a PilA-N-like protein or a heterodimer, or whether a cytochrome was measured instead of any pilus whatsoever is not known. However, it is interesting that Malvankar and co-workers reported using the *Aro5* strain to produce OmcS filaments,[13] begging the question of whether the putative Aro5 pilus was really the OmcS filament.

(13) Yalcin, S. E.; O'Brien, J. P.; Gu, Y.; Reiss, K.; Yi, S. M.; Jain, R.; Srikanth, V.; Dahl, P. J.; Huynh, W.; Vu, D.; et al. Electric field stimulates production of highly conductive microbial

OmcZ nanowires. *Nat Chem Biol* **2020**, *16* (10), 1136-1142. DOI: 10.1038/s41589-020-0623-9 From NLM Medline.
